# Structural and oligomeric characterization of substrate- and product-selective nylon hydrolases

**DOI:** 10.64898/2026.03.11.711162

**Authors:** Nikolas Capra, Célestin Bourgery, Jerry M. Parks, Dana L. Carper, John F. Cahill, Joshua K. Michener, Flora Meilleur

**Author notes:** Corresponding authors: N.C. and F.M. Notice: This manuscript has been authored by UT-Battelle, LLC under Contract No. DE-AC05-00OR22725 with the U.S. Department of Energy. The United States Government retains and the publisher, by accepting the article for publication, acknowledges that the United States Government retains a non-exclusive, paid-up, irrevocable, world-wide license to publish or reproduce the published form of this manuscript, or allow others to do so, for United States Government purposes. DOE will provide public access to these results of federally sponsored research in accordance with the DOE Public Access Plan (http://energy.gov/downloads/doe-public-access-plan).

## Abstract

Enzymatic degradation of synthetic polymers has attracted broad interest because it offers environmental and manufacturing advantages compared to traditional mechanical and chemical breakdown approaches. Enzymes are highly specific and reaction conditions are generally aqueous and require low pressure and temperature, resulting in lower energy consumption and lower chemical waste production. Here we report the biochemical and structural characterization of three newly discovered enzymes capable of nylon hydrolysis: Nyl10, Nyl12 and Nyl50. Using solution characterization techniques, we found that the enzymes adopt a single oligomeric state consistent with a tetramer over a wide range of concentrations. X-ray crystallographic structures of all three enzymes support the association into tetramers. Comparison of ligand-bound X-ray crystal structures of Nyl10 and Nyl12 with the previously determined structure of Nyl50 identified key structural determinants involved in ligand binding. Noticeably, a flexible loop found in several polyamide degrading enzymes is observed to flip towards (closed conformation) and away (open conformation) from the active site upon ligand binding. Analysis of adduct and surrogate substrate-bound enzyme complex structures provide a model for substrate binding directionality. Finally, activity assays showed that both Nyl10 and Nyl12 can hydrolyze ester bonds, and that Nyl12 has the highest activity toward PA66, identifying it as the best candidate for protein engineering for efficient nylon hydrolysis.

## Introduction

Nylons are synthetic polyamides that are widely used in everyday products (e.g., automotive, textiles), due to their high strength, elasticity, as well as abrasion and chemical or thermal resistance. Nylon 6 (polyamide 6 or PA6) and nylon 6,6 (polyamide 6 or PA66) represent the most common nylon variants and differ in molecular structure and synthesis. Specifically, nylon 6 is obtained when the ω-amino acid 6-aminocaproic acid (6-ACA) or its lactam (ε-caprolactam) undergoes self-condensation, while nylon 6,6 requires adipic acid and hexamethylenediamine.^1^ The energy required for the synthesis of nylon precursors and the consequent generation of greenhouse gasses are considerable.^2^ Increasing energy costs and concerns on environmental impact, in concert with a constant increase in demand and more attention to sustainability, have underlined the urgency of recycling and the design of a circular economy of plastics, including polyamides. However, polyamide recycling remains challenging. The most common recycling methods for plastic waste are mechanical processing (e.g. melting and shredding), that alters the properties of the nylon fibers and shortens the lifespan of the materials^3–5^, and chemical processing, that requires high temperatures and strong acids resulting in the generation of caustic waste and undesired side products.^6,7^

Several studies on polyethylene terephthalate (PET) enzymatic degradation have demonstrated that engineered PETases are capable of hydrolyzing PET into monomers from low-purity feedstocks which can then be chemically repolymerized into high-quality polymers^8–14^. Biocatalytic yields obtained for PET have opened the door to the valorization of more recalcitrant polymers, such as nylon. We have recently shown that the ability of nylon-active enzymes to generate oligomeric hydrolysis products results in a higher degree of polymerization when these hydrolysis products are used instead of small-molecule salts (i.e., adipic acid and diamine).^15^ However, efficient enzymatic degradation of polyamides remains a challenge due to their physical and chemical properties including, but not limited to substrate crystallinity and high molecular weight (MW), and to the overall low activity of nylon hydrolases identified to date.^15–17^

The first nylon 6 oligomer hydrolase, NylC, was the focus of enzyme engineering campaigns after being extensively characterized by x-ray crystallography and mutational analyses, including a study reporting the oligomeric assembly model.^18,16,17,19,20^ The model proposed that a functional tetramer assembled via a specific assembly sequence and that the oligomeric state populations are dynamic and influenced by mutations. Recently, we reported the identification of several thermostable, product and substrate selective nylon hydrolases, active toward the most common nylon variants (PA6 and PA66), using a diversity panel and systematic analysis of product formation. Out of 95 discovered enzymes, three (named Nyl10, Nyl12 and Nyl50) showed different substrate specificity for either PA6 or PA66 oligomers and different product distribution.^15^ The crystal structure determination of the PA66-selective Nyl50 allowed the identification of a putative substrate access tunnel thanks to PEG molecules bound in proximity of the catalytic residue.^15^ The mechanisms involved in substrate and product selectivity, and the role of oligomerization in catalysis remain unclear.

**Scheme 1.**
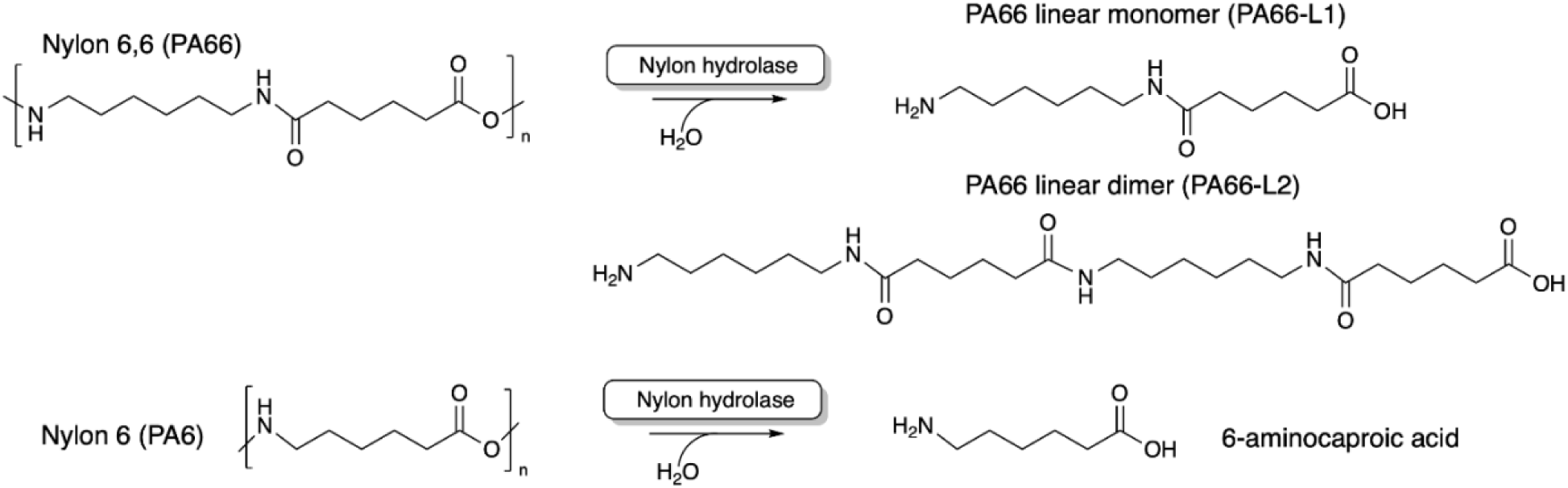
Schematic representation of the nylon hydrolysis reaction for PA66, PA6 and their products.

In this work, we conducted structural studies using X-ray crystallography and biophysical analyses on the recently identified nylon hydrolases with different substrate and product specificities, Nyl10, Nyl12 and Nyl50, to elucidate sequence-structure-function relationships and understand the role of oligomerization in the functional enzymes. We report the first crystal structures of Nyl10 and Nyl12, along with ligand-bound structures of the previously characterized Nyl50. Based on crystallographic analysis and computational modeling, we annotate the architecture of the access tunnel with respect to substrate binding directionality. Finally, we investigate the oligomerization states of each enzyme in solution under different conditions. These studies revealed that nylon hydrolases form stable tetramers over a wide range of concentrations and that key structural determinants involved in enzymatic activity, including a highly flexible loop, could be the target of future enzyme engineering campaigns.

## Materials and Methods

### Protein expression and purification

Reagents and supplies for enzyme expression and purification were purchased from VWR, FisherScientific and SigmaAldrich. Expression and purification protocols were adapted from Drufva et al.^15^ Briefly, plasmids containing the genes for Nyl50, Nyl10 and Nyl12 in vector pET28a(+) were transformed into *E. coli* BL21(DE3) competent cells (New England Biolabs, USA), plated on lysogeny broth (LB) agar plates supplemented with 50 μg/mL kanamycin and incubated at 37°C overnight. A single colony was inoculated into 8 mL of LB supplemented with 50 μg/mL kanamycin and incubated at 37°C overnight. The next day, 5 mL of pre-culture were transferred into 2.8 L baffled flasks containing 500 mL of terrific broth (TB) supplemented with 50 μg/mL kanamycin. Cells were grown at 37°C until OD_600_ reached 1.2-1.4. Prior to induction, cell cultures were cooled down to 4°C for 15 minutes. Protein expression was induced by adding of isopropyl β-D-1-thiogalactopiranoside (IPTG) to a final concentration of 0.25 mM and incubated at 20°C overnight (19h), shaking at 110 rpm. Cells were harvested by centrifugation at 4000 *x g* at 4°C. Cell pellet was resuspended in 35 mL of 20 mM Tris buffer pH 7.9 supplemented with 1 mg/mL lysozyme and 1 cOmplete Protease Inhibitor Cocktail tablet (Roche) and sonicated in ice for 15 minutes with 5 seconds intervals. As a first purification step, the lysate was subjected to heat-treatment by incubating it at 65°C for 1 hour and successively centrifuged at 25000 *x g* at 4°C for 30 minutes. Supernatant was then recovered and loaded into a Q Trap HP 5 mL column (Cytiva, Sweden) equilibrated with 20 mM Tris buffer pH 7.9 using an ÄKTA Pure FPLC system (Cytiva, Sweden). Elution was carried out using a stepwise method with buffer containing 20 mM Tris pH 7.9, 1 M NaCl. Nyl50 eluted at 200 mM NaCl in two close peaks, while Nyl10 eluted at 300 mM in a single peak. Pooled fractions were concentrated to a final volume of 5 mL and loaded into an HiLoad 16/600 Superdex 200 pg size exclusion column (SEC) (Cytiva, Sweden) equilibrated with 20 mM Tris buffer pH 7.9, 100 mM NaCl. Conversely, Nyl12 did not bind the Q Trap column, due to its higher pI, thus the flow-through was directly loaded onto the size exclusion column. All three proteins eluted in single sharp peak, with a purity of at least 95%, as assessed by SDS-PAGE. Finally, fractions containing the target protein were pooled and concentrated using a Vivaspin®20 10K polyethersulfone filter unit (Sartorius) to about 10 mg/ml, flash frozen in LN_2_ and stored at −20°C. BioRad gel filtration standards (BioRad, USA) were used to generate a calibration curve for approximate molecular weight calculation. Analytical SEC was carried out using 50 μL of purified samples using an ÄKTA Pure fitted with a Micro kit and a Superdex 200 Increase 3.2/300 at 0.05 mL/min.

### Crystallization and structure determination

Crystallization screening of Nyl10 and Nyl12 was conducted at 20°C at the National High-Throughput Crystallization Screening Center at the Hauptman–Woodward Medical Research Institute (Buffalo, NY, USA).^21,22^ Several conditions gave small, three-dimensional crystals for both proteins and were reproduced successfully. For Nyl10 the condition from HR Slice pH B2 (0.5 M DL-malic acid pH 4.7, 15% w/v PEG 3350) produced the best diffracting crystals. Greiner 72-well micro-batch plates were used to set up micro-batch drops under oil at different protein:cocktail ratios (1:2, 1:1, and 2:1). Protein solution of Nyl10 at about 7 mg/mL was mixed with the crystallization cocktail up to a final volume ranging from 2 to 4 μL and subsequently overlaid with paraffin oil. Various small, three-dimensional crystals appeared within 24-48 hours, with the best diffracting crystals grown in a 1:1 ratio. Prior to data collection, crystals were briefly soaked in a cryoprotectant solution containing the crystallization cocktail and 20% v/v glycerol and flash-frozen in LN_2_. Attempts to obtain a ligand-bound structure including several experiments of co-crystallization and soaks with PA66 rinsate were unsuccessful. A suitable x-ray diffraction dataset to 1.3 Å was collected at 100 K at beamline 12-1 at the Stanford Synchrotron Radiation Light (SSRL) in Stanford, California, USA. Diffraction data were indexed and integrated in the space group C 2 using XDS^23^ from autoPROC^24^ installed in the beamline data processing pipeline, then scaled and merged with Aimless^25^ from the CCP4 suite.^26^ The asymmetric unit was estimated to contain six protein molecules with a Matthews coefficient of 2.13 and a solvent content of 42%. Molecular replacement was carried out using Phaser^27^, using an AlphaFold2 (AF2) model as a search model.^28^ Next, a few cycles of refinement using REFMAC5^29^, alternated by model building and water placement using Coot^30^, were sufficient to complete the structure.

Similarly, Nyl12 crystallization screening produced crystals in several conditions containing HR Ionic Liquid Screen additives (Hampton Research, USA). Micro-batch crystallization experiments were set up as described above. Single, three-dimensional crystals appeared within 24-48 hours in conditions containing 0.09 M 4-(2-hydroxyethyl)-1-piperazine ethanesulfonic acid (HEPES) pH 6.8, 27% w/v PEG 3350 and different ionic liquids. The best diffracting crystals were observed in drops containing 0.09 M HEPES pH 6.8, 27% w/v PEG 3350 and 5% v/v choline acetate (HR Ionic Liquid 5) in a protein:cocktail ratio of 2:1. A MiTeGen MicroRT system (MiTeGen, USA) was used to mount a crystal for room temperature data collection on a Rigaku HighFlux HomeLab instrument equipped with a MicroMax-007 HF X-ray generator and Osmic VariMax optics. A dataset to 1.9 Å resolution was recorded using an Eiger 4 M hybrid photon counting detector. Diffraction data were indexed and integrated using the CrysAlis Pro software suite (Rigaku, USA) in space group P 2 2_1_ 2_1_, scaled and merged with Aimless. A search for two copies of the Nyl12 monomer in the asymmetric unit was run, as suggested by a Matthews coefficient of 2.39 and a solvent content of 48.5%, using an AF2 model as a search model. Subsequently, a few rounds of refinement using REFMAC5, alternated by model building with Coot were sufficient to complete the structure. Additionally, attempts to grow crystals of sufficient size for neutron diffraction yielded large single crystals. The experiment was carried out in a 1.5 mL Eppendorf tube by adding 20 μL of silica gel (Hampton Research, USA), obtained by adding sodium silicate to acetic acid in a 1:1 ratio and allowing for the gel to polymerize and cure for about an hour. Excess water produced by the polymerization reaction was dried with a paper wick. Next, 20 μL of protein solution was added to the gel and allowed to cure for about an hour, followed by the addition of 40 μL of crystallization cocktail. The tube was sealed with parafilm and incubated at 18°C. A single crystal of about 0.7 x 0.6 x 0.4 mm was observed after 4 weeks. Prior to data collection, the crystal was briefly soaked in a cryoprotectant solution containing the crystallization cocktail and 25% v/v glycerol. An X-ray diffraction dataset to 1.75 Å was collected at 100K at beamline 12-1 at the SSRL. Diffraction data were indexed and integrated in the space group P 2_1_ 2_1_ 2_1_ using XDS^23^ from autoPROC^24^, then scaled and merged with Aimless^25^ from the CCP4 suite.^26^ The asymmetric unit was estimated to contain between six and eight protein molecules. Molecular replacement was carried out using Phaser and a monomer of the previous determined structure was used as a search model, with the best solution found for eight molecules (Matthews coefficient 2.06, solvent content 40.3%). Structure refinement was carried out using REFMAC5^29^ and manual model building using Coot.^30^

Crystals of Nyl50 were grown as previously described.^15^ Ligand-bound structure of Nyl50 were obtained by soaking the crystals in a drop containing the crystallization cocktail supplemented with 5 mM of N-(4-nitrophenyl) butanamide (N4NB) and 4.5 mg/mL of chemically synthesized PA66 linear monomer (PA66-L1) solution. Both N4NB and PA66-L1 stock solutions were obtained by resuspending the compounds in the protein storage buffer. Crystals were soaked in a 5 μL drop between 1, 18, and 40 hours at room temperature. Diffraction datasets to 2.0 and 3.1 Å resolution were recorded for butyrate-bound and acyl-enzyme, respectively, at room temperature, using the in-house X-ray source mentioned above. Diffraction data were indexed and integrated using the CrysAlis Pro software suite (Rigaku, USA) in space group P 2 2_1_ 2_1_, scaled and merged with Aimless. The crystals belonged to the same space group of the previous Nyl50 structure (PDB ID 9DYS) with nearly identical unit cell dimensions, allowing for the previously determined structure to be used for initial refinement and electron-density map calculations. Coordinates for PA66-L1 were generated with Avogadro^31^ and energy minimization was carried out using the MMFF94 force field.^32^ The constraints file was generated using eLBOW^33^ from the Phenix^34^ suite of programs. The butyrate molecule (Chem ID BUA) was imported from the PDB dictionary. Several cycles of REFMAC5^29^ alternated with manual model building with Coot^30^ were performed to complete the structures. Data collection and refinement statistics are shown in Table S1.

### Colorimetric activity assays

Amide hydrolytic activity was measured against chromogenic substrates N-(4-nitrophenyl) butanamide (N4NB), N-(4-aminophenyl) butanamide (N4AB), and N-(4-nitrophenyl) butyrate (N4NB), adapting the activity screening protocol reported by Bell et al.^19^ Reactions were carried out at 200 μL scale in microtiter plates using 500 μM of either N4NB or N4AB, dissolved in DMSO, in reaction buffer (20 mM Tris, pH 7.9, 100 mM NaCl) and the reaction was initiated by adding 3 μM of enzyme. The plate was sealed with optically clear film and the release of *p*-nitroaniline (*p-*NA), *p*-phenylenediamine (*p-*PD), and *p*-nitrophenol (*p-*NP) were monitored by change in absorbance at 410, 286, and 405 nm, respectively, using a Molecular Devices SpectraMax i3 over the course of 24 hours at 30°C with measurements every 10 minutes. Additionally, we carried out spectra measurements on the N4AB-incubated samples after the reaction ended, to ensure that the formation of *p-*PD was not overlooked due to the fixed wavelengths monitored. For each experiment, negative control samples containing either no enzyme or no substrate were included and reactions were carried out in triplicates.

### Computational modeling

Enzyme-substrate models of Nyl12 with PA66-L2 were generated as described previously.^35^ Briefly, Boltz-2^36^ and the ColabFold multiple sequence alignment server^37^ were used to generate 15 models of the complex, which then underwent energy minimization with OpenMM 8.0^38^ using the Amber FF19SB force field^39^ for the protein and General Amber Force Field (GAFF)^40^ for the substrate. PyRosetta 4.0^41,42^ was used to calculate enzyme-substrate interaction energies (i.e., *interface_delta*). Models filtered based on their active site geometries, Boltz-2 confidence scores, and interaction energies. The Protein-Ligand Interaction Profiler (PLIP)^43^ was used to identify polar and nonpolar interactions between the enzyme and substrate in the models.

### Dynamic light scattering

Dynamic light scattering (DLS) experiments were performed on a Wyatt Technologies DynaPro™ NanoStar™ I instrument. The instrument was equilibrated to 20°C and 5 μL of sample were loaded into a Wyatt disposable MicroCuvette. The instrument was set to 10 acquisitions with 5 seconds as the acquisition time. Prior to data acquisition, the instrument was allowed to re-equilibrate to 20°C after sample loading. Experiments were performed at least in duplicate.

### SAXS data acquisition and analysis

SAXS was performed in-house using a Rigaku HighFlux HomeLab instrument equipped with a MicroMax-007 HF X-ray generator coupled with Rigaku BioSAXS 2000 mounting OptiSAXS CMF focusing optic. Prior to data collection, the instrument was calibrated using silver behenate for a configuration with minimum *q* of ∼0.015 Å^−1^. A concentration range (1.0, 2.5, 5.0, 10 mg/mL) was explored for each enzyme, generating four datasets per enzyme with 180-minute exposure for each sample. The measured raw data were buffer subtracted using the SAXSLab software package (Rigaku) and analyzed using the RAW program from the BioXTAS software package.^44^ Guinier analyses and molecular weight calculations were carried out using the RAW program. Indirect Fourier Transform (IFT) and pair distance distribution analyses (P_r_) were calculated using GNOM^45^, from the ATSAS^46^ software package, and BIFT via the RAW program interface. SAXS profiles of Nyl10, Nyl12 and Nyl50 crystal structures were computed and fitted to SAXS experimental data using the webserver FoXS^47,48^ and the oligomeric distribution for the dimer and the tetrameric structures were computed using OLIGOMER^49^ from the ATSAS online webserver.

### Analytical ultracentrifugation

The analytical ultracentrifugation experiments (AUC) for Nyl50, Nyl10 and Nyl12 were performed using previously published protocols for NylC-GYAQ.^16^ Briefly, all three enzymes were analyzed by sedimentation velocity at 20°C in 20 mM Tris HCl, pH 7.9, and 100 mM NaCl using a Beckman-Coulter Optima XL-A analytical ultracentrifuge (Beckman-Coulter, USA) with the rotor pre-equilibrated at 3,000 rpm for 3 hours to prevent any thermal discrepancies. Centrifugation was performed at 40,000 rpm by monitoring A_280_ at 5 min intervals. For sample loading, all experiments used double-sector 12-mm-thick charcoal-epon centerpieces and matched quartz windows. Buffer density, viscosity and partial specific volume (Vbar) were calculated using SEDNTERP3^50^ and successively the experiments were analyzed using SEDFIT18.1.^51,52^

### Mass photometry

Mass photometry (MP) measurements were carried out using the Refeyn TwoMP mass photometry setup and measurements were recorded using the Refeyn AcquireMP software. To set the experimental stage, a 6-well silicon gasket was positioned onto a clean microscope coverslip, which was then inserted in the instrument and locked in place by two magnets. Mass calibration was performed using protein standards of known masses, specifically thyroglobulin (660 kDa) and bovine albumin serum (BSA, 66 kDa). All three nylon hydrolase samples were diluted to a concentration of 200 nM as an initial stock. To perform the measurement, 18 μL of filtered buffer was added in the designated well and the focus was found using the droplet dilution setting in the AcquireMP software. After achieving focus, 2 μL of protein sample were added to the buffer droplet, to a final concentration of 20 nM, and gently mixed three times. Then, data collection was initiated and recorded for one minute. Subsequent data processing and analysis were conducted using Refeyn DiscoverMP software.

### Steady-state kinetics

Michaelis-Menten kinetics were employed to evaluate the depolymerization efficiency of Nyl50, Nyl10, and Nyl12. The protocol was adapted from Puetz et al.^53^ A fixed concentration of enzymes (500 nM) was tested with varying substrate concentration (38 to 700 g·L⁻¹) of PA66 powder (particles sizes < 300 μM), which was prepared according to the previously reported method.^54^ The dried powders were washed with methanol to remove unreacted precursors and byproducts.^15,54^ Depolymerization reactions were conducted by incubating the mixtures at 75 °C without shaking for 2 hours in 200 mM Tris buffer, pH 8. Hydrolysis products were quantified using our established I.DOT/OPSI-MS analytical methods.^15^ Initial velocities (v, μM·s⁻¹) were plotted as a function of substrate concentration ([S], g/L) in GraphPad Prism (version 10.6.1 (799)). Data were fitted to the standard Michaelis–Menten equation using nonlinear regression:

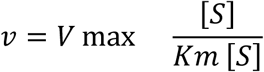

All experiments were performed in biological triplicate, and values are reported as mean ± standard deviation.

### I.DOT/OPSI-MS analysis of enzyme activity

The immediate drop-on-demand technology (Dispendix GmbH, Stuttgart, Germany) coupled with open port sampling interface mass spectrometry (I.DOT/OPSI-MS) was used to characterize and quantify PA hydrolysis products.^55^ Reactions were diluted 1:100 (v/v) in HPLC grade water having 500 nM propranolol. Propranolol intensities were used to normalize droplet-to-droplet variability. 40 μL of diluted reactions were transferred to I.DOT S.100 96-well plates and analyzed immediately by ejecting 20 nL of sample into the OPSI-MS, having a 170 μL/min flow of 75/25/0.1 (v/v/v) acetonitrile/water/formic acid. The Sciex 7500 triple quadrupole mass spectrometer (SCIEX, Concord, Ontario, Canada) was operated in positive ion mode using the following parameters: gas setting 1 = 90, gas setting 2 = 60, electrospray voltage = 5.5 kV, capillary temperature = 200 °C, and dwell time = 20 ms. Multiple reaction monitoring scans were 260.1→183.0; collision energy (CE)=26 eV (propranolol), 245.19→100.11; CE=27 eV (PA66-linear monomer, diad), 471.35→100.11; CE=65 eV (PA66-linear dimer, tetrad), 245.19→114.09; CE=26 eV (PA6-linear dimer), 358.27→114.09; CE=50 eV (PA6-linear trimer), and 471.35→114.09; CE=65 eV (PA6-linear tetramer). Measurements of PA66 and PA6 oligomers were always collected in separate experiments. Each sample was averaged over three technical replicates. Reported concentrations are the mean of biological replicates. Each oligomer was quantified using a 12-point calibration curve (0-50 μg/mL) from respective standards. Analysis was performed using custom software (ORNL I.DOT-MS Coupler v2.50 and PyFindPeaks).

## Results and Discussion

### Structural analysis of Nyl10 and Nyl12 structures

Nyl10, Nyl12, and Nyl50 are part of the NylC superfamily but share relatively low sequence identity (>40%, Figure S1). Different substrate and product specificity was reported for these enzymes. Nyl10 and Nyl50 selectively hydrolyzed PA66 while Nyl12 hydrolyzed both PA6 and PA66. Moreover, Nyl12 hydrolyzed PA6 mostly into linear trimeric and tetrameric products. Nyl10 and Nyl50 generated mostly PA66 linear dimeric products, whereas NylC yielded predominantly diads from both substrates.^15^ Thus, to understand the structural differences and identify the structural determinants of substrate and product selectivity in these enzymes, we crystallized and determined a 1.3 Å resolution X-ray crystallographic structure of Nyl10 (PDB ID 9YYS) at 100 K and two structures of Nyl12 at 1.75 Å (PDB ID 9Z0B) and 1.9 Å (PDB ID 9Z0A) at 100 K and room temperature, respectively. For clarity, we refer to the Nyl12 structures as Nyl12-CT and Nyl12-RT. For both Nyl10 and Nyl12 structures, the monomers contained an α and a β subunit, similar to the to the previously described Nyl50.^15^

For Nyl10, the asymmetric unit contained six monomers, arranged in a tetramer and a dimer. The latter showed an A/D dimer assembly, arranged in an αββα structure, and formed a tetramer around the crystallographic 2-fold symmetry axis. The A/B/C/D tetramer observed (Figure 1A) is similar to the one described for NylC from *Agromyces sp.*^18^, with a Cα backbone RMSD of 1.15 Å, indicating that no major differences in secondary structure are present despite the low sequence identity (23%). Electron density was absent for residues 201-210 located at the C-terminal domain of the α subunit, as a result of the autocatalytic cleavage that occurs to release the catalytic threonine, Thr211, located at the N-terminus of the β subunit. Absence of electron density for the residues preceding the catalytic threonine suggests that this tail region becomes highly flexible upon its release, consistent with the previously described Nyl50 and NylC.^15,16,18^

**Figure 1.**
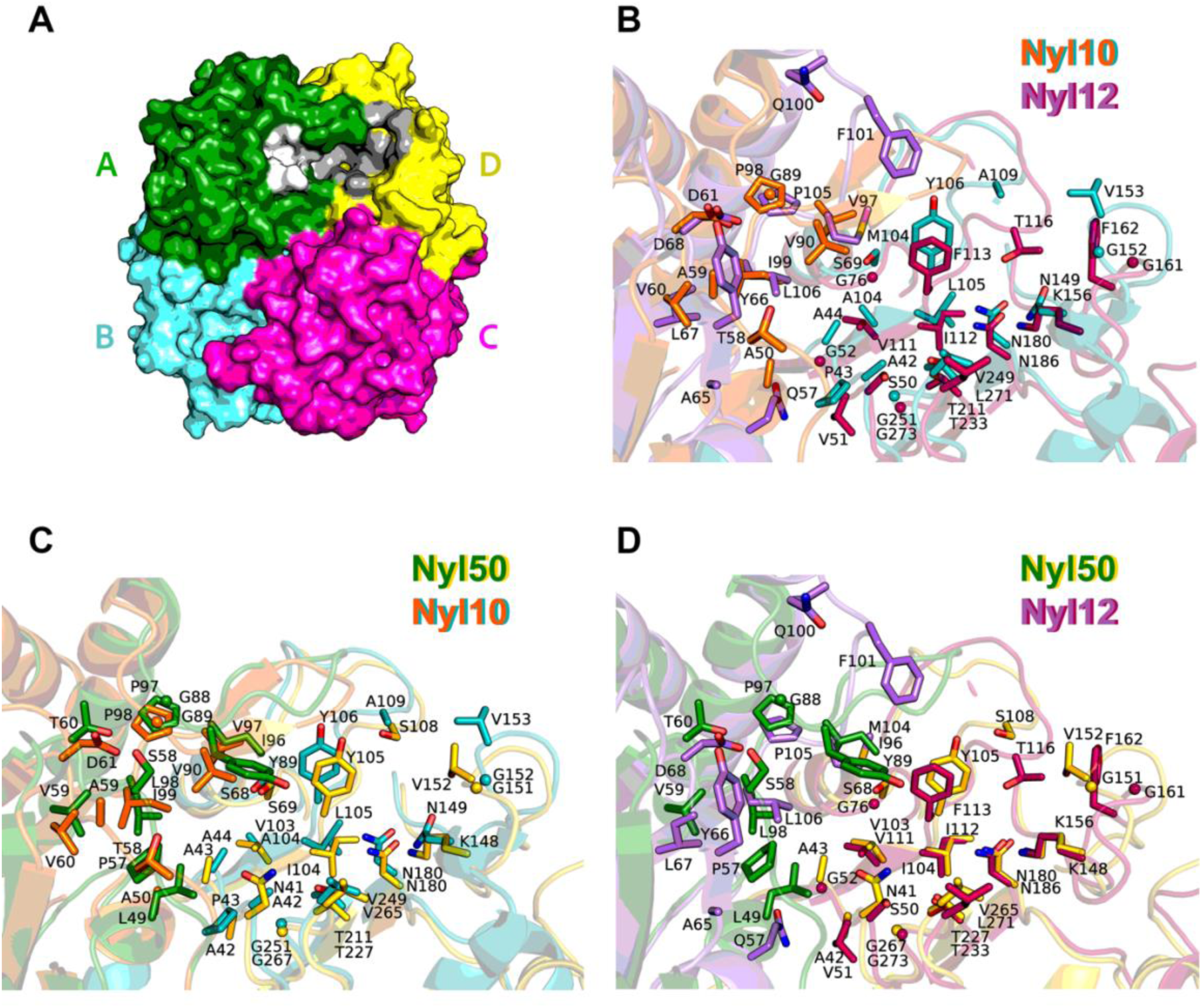
Nylon hydrolase active sites. **A**. Tetrameric assembly of a nylon hydrolase represented as a surface rendering colored by chain. Patches indicating the location of residues that form the active site access tunnel, shown only in molecules A and D for clarity, are colored white and grey, respectively. **B-D**. Comparison of active site residues at the A/D dimer interface (cartoon, bicolor indicating the different monomers), in **B.** Nyl10 (orange and teal) and Nyl12 (purple and maroon). **C**. Nyl10 (orange and teal) and Nyl50 (forest green and yellow). **D**. Nyl12 (purple and maroon) and Nyl50 (forest green and yellow). The active site of Nyl10 is the most similar to Nyl50. No major differences in secondary structures are present, except for the loop harboring Phe101 in Nyl12.

The Nyl12-RT data revealed a dimer in the asymmetric unit, but in the A/B assembly, forming a tetramer around the crystallographic 2-fold symmetry axis. Monomer A of Nyl12-RT superimposed onto monomer A of the Nyl50 structure (PDB ID 9DYS) with a Cα backbone RMSD of 1.0 Å, indicating the conservation of the overall structure fold. Consistent with the other nylon hydrolases, Nyl12-RT also lacks electron density for residues 228-232, which belong to the highly flexible C-terminal tail of the α subunit preceding the catalytic residue, Thr233. The crystal structure of Nyl12-CT was determined in space group P 2_1_ 2_1_ 2_1_, different from the P 2 2_1_ 2_1_ of Nyl12-RT, with eight molecules in the asymmetric unit assembled in a A/B/C:A/B:C/D:D pattern. We hypothesize that the difference in space group is due to the different type of crystallization experiment. The crystal of Nyl12-RT was obtained by microbatch-under-oil experiments, with a full growth time of ∼48 hours, while the Nyl12-CT crystal was obtained by liquid/gel diffusion using silica hydrogel, with a full growth time of ∼3 weeks. We hypothesize that the slower growth rate affected the crystal packing, resulting in a different crystal morphology (Figure S2) and thus in a different space group. Despite the cryogenic temperature, the different space group, and the higher resolution, the tail remains partially unsolved and there are no differences in the overall structure compared with Nyl12-RT, as indicated by the Cα backbone RMSD of 0.4 Å. Gratifyingly, we fully modeled a loop located near the access tunnel at the A/D dimer interface comprising residues 95-107 in chains A, G, and H, which is otherwise partially unsolved due to its flexibility.

### Active sites of Nyl10 and Nyl12

The conserved overall fold reflects on the conserved shape of the tunnel-like active site in both Nyl10 and Nyl12, compared to Nyl50. The structural elements in the active site appear to be highly conserved in terms of steric hindrance, charge, and orientation. Specifically, considering the functional dimer A/D, with the catalytic Thr in monomer D, the bulk of the active site is located in monomer D and shows the most conservation, while the residues from monomer A contributing to the active site show more variability and are located on a loop at the dimer interface that appears to be highly flexible (Figure 1). A comprehensive list of the residues forming the access tunnel and active site is given in Table S2.

Interestingly, comparison with Nyl50 revealed that between Nyl10 and Nyl12, the active site of Nyl10 is the most similar to Nyl50, except for residue Asn149 that is a conserved lysine in Nyl50, Nyl12, and NylC (Lys148, Lys156, and Lys189, respectively). Moreover, Nyl12 presents bulky or polar residues, such as Gln57, Tyr66 and Met104, whereas Nyl50 and Nyl10 have shorter and apolar residues such as Leu49 and Ala50, Ser58 and Ala59, Ile96 and Val97, respectively. The pocket volumes were 631 Å^3^ for Nyl50, 777 Å^3^ for Nyl10, and 706 Å^3^ for Nyl12. It has been observed that steric hindrance, hydrophobicity and the equilibrium between charged residues in the active site can substantially affect substrate binding and the overall amide-hydrolytic activity.^56^ These differences may explain the different substrate preference between Nyl12 and Nyl10, the latter of which hydrolyzes PA66 in a comparable way to Nyl50.

Although the enzymes were crystallized in the absence of a nylon-derived substrate, we observed non-protein electron density in the active site of the Nyl12 structures. The Nyl12-RT structure showed an elongated F_O_-F_C_ electron density at about 2.5 Å and a shorter density at ∼4.2 Å from the catalytic Thr233 in monomers A and B, respectively. Considering the crystallization conditions, we modeled triethylene glycol (Chem ID PGE) in chain A, while the density in chain B was consistent with the shorter di(hydroxyethyl)ether (PEG). We note that PEG molecules from the crystallization buffer were previously observed in the active site of Nyl50.^15^ The crystal structure of Nyl12-CT showed different non-protein densities in the active sites that were consistent with the crystallization buffer components choline (CHT) and acetate (ACT) in monomer A, with a HEPES and a glycerol (GOL) molecule in monomer C, a PEG and a CHT molecule in monomer D, a single HEPES molecule in monomers E and F, and an ACT molecule in monomer H. Notably, although the PEG molecule is bound in the active site close to residue Thr233, the choline molecules are buried at the interface portion of the tunnel and the HEPES molecule is located closer to the surface of the protein at about 8 Å from the O_γ_ of Thr233, interacting with the sidechain of residue Leu299 from monomer C via its piperazine ring and via hydrophobic interaction between the Cδ of Ile295 and its C7, as well as with Phe113 and with the side chain of residue Arg159 in monomer D via its sulfonate group (Figure 2).

**Figure 2.**
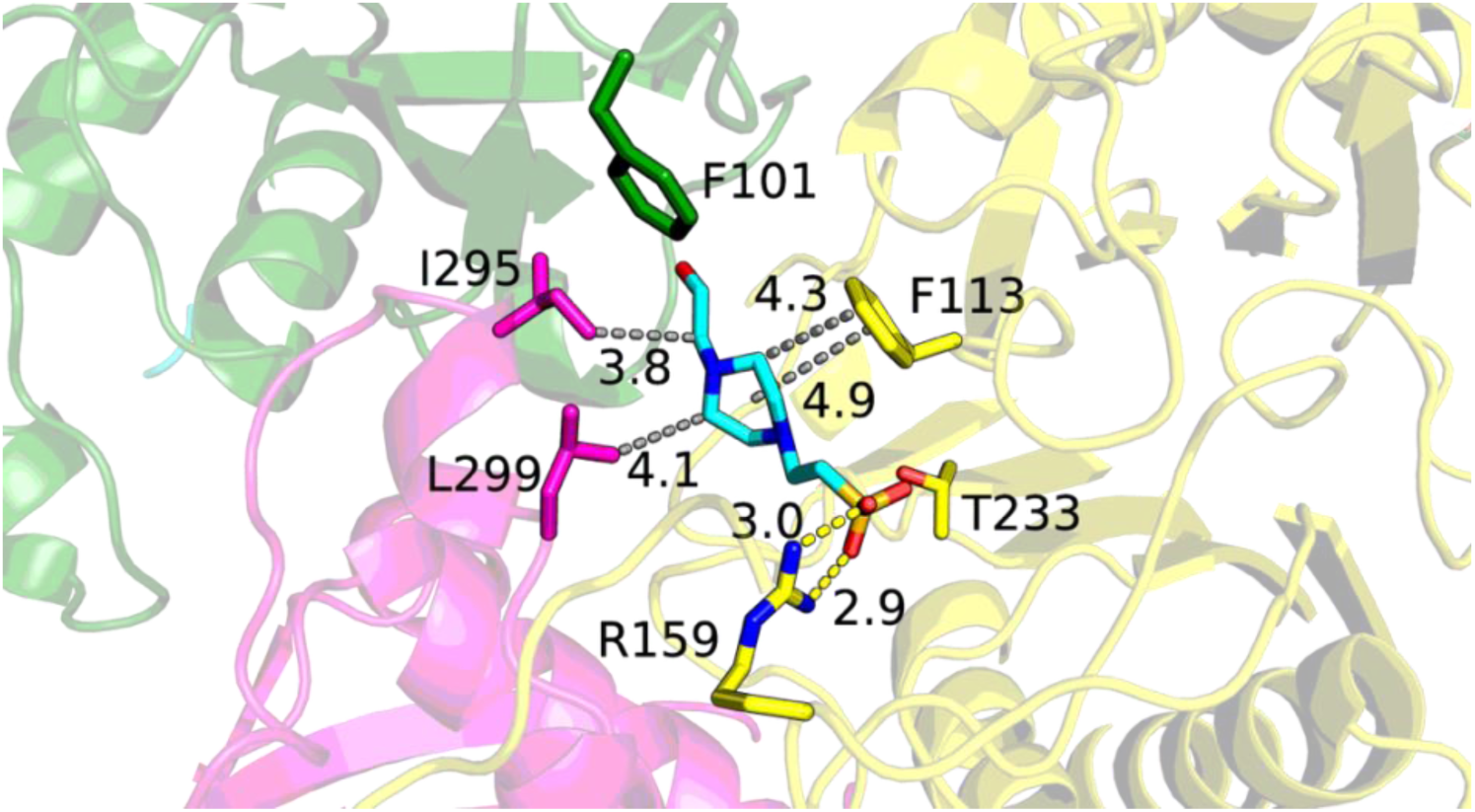
Nyl12 with a HEPES molecule bound in the active site. Residues Ile295 and Leu299 stabilize the ligand via hydrophobic interactions, suggesting that the active site is not limited to the A/D dimer, but a third subunit may contribute to its architecture. Arg159 interacts with HEPES via polar interactions. Color scheme is the same as in Figure 1: A/D dimer in green/yellow and chain C in magenta. Distance between Phe113 and HEPES is measured between ring centroids. Hydrophobic contacts (grey) and hydrogen bonds (yellow) are shown as dashed lines. Distances are in Å.

Similarly, both the acetate molecules interact with the side chain of Arg159, which is oriented toward the active site. Other members of the Ntn-hydrolase family present an arginine residue in proximity of the active site and exposed to the solvent, which in the conjugated bile amide hydrolase (CBAH) was deemed responsible for facilitating the expulsion of the hydrolysis product.^57^ Comparison between Nyl12-RT and Nyl12-CT structures showed a rotation of the side chain of Arg159 of ∼180°, pointing outward toward the solvent in Nyl12-RT due to the absence of a negatively charged ligand (Figure 3A). Localized at a similar position in Nyl50 is Arg153 and Arg156 in Nyl10. These conserved arginine residues correspond topologically to Arg330 in NylC, which is putatively responsible for stabilizing the substrate.^20^ The position and the 180° side chain flip suggests that this Arg residue may act as a switch, stabilizing the substrate and ushering the reaction products out of the active site tunnel. Interestingly, Arg330 in NylC belongs to monomer C, while in Nyl50, Nyl10, and Nyl12 the topological analogues pertain to the same monomer that hosts the bulk of the active site (e.g., monomer D in the A/D dimer) (Figure 3B) suggesting that, while the dimer hosts the bulk of the active site, a third subunit may contribute to its architecture.

**Figure 3.**
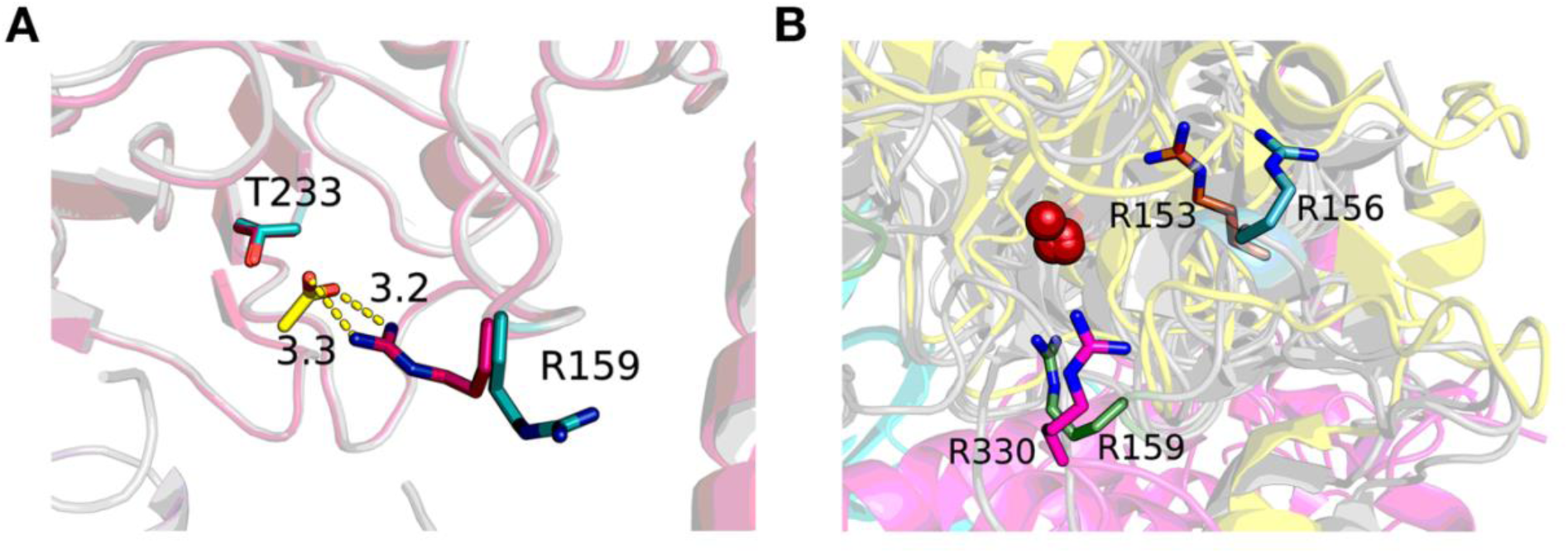
**A.** Arg159 in Nyl12 switches from a solvent-oriented position (teal) to an active site-oriented position (hot pink) position in the presence of a ligand with a negatively charged group (ACT, yellow). **B.** Superposition of Nyl10, Nyl12, and Nyl50 A/D dimers (grey) and NylC tetramer (colored) showing spatial localization of the conserved Arg156 (teal), Arg153 (orange) and Arg159 (green) in Nyl10, Nyl50, and Nyl12, respectively, on chain D (in yellow for NylC) compared to the analogous Arg330 on chain C in NylC (magenta). For reference, the catalytic threonine is represented as red spheres.

### Analysis of loop at the tunnel access site

The flexible loop at the A/D dimer interface tunnel access site (residues 95-107, the AS-loop hereafter) observed in monomers A, G, and H in the Nyl12-CT structure was trapped in different conformations, with no significant deviation of the surrounding secondary structure elements to which it is anchored. The three different loop conformations suggest an oscillation from a position close to monomer A (A-swing) to a position closer to monomer D (D-swing) with a maximum distance of 7.7 Å between the respective Phe101 Cα atom coordinates and an intermediate position on the same axis (Figure 4A). Comparison with Nyl12-RT revealed that the loop shifted toward the active site (closed conformation), positioning the Phe101 Cα at ∼9.6 Å and 7.1 Å vertically compared to the A-swing and D-swing orientation, respectively (Figure 4B).

**Figure 4.**
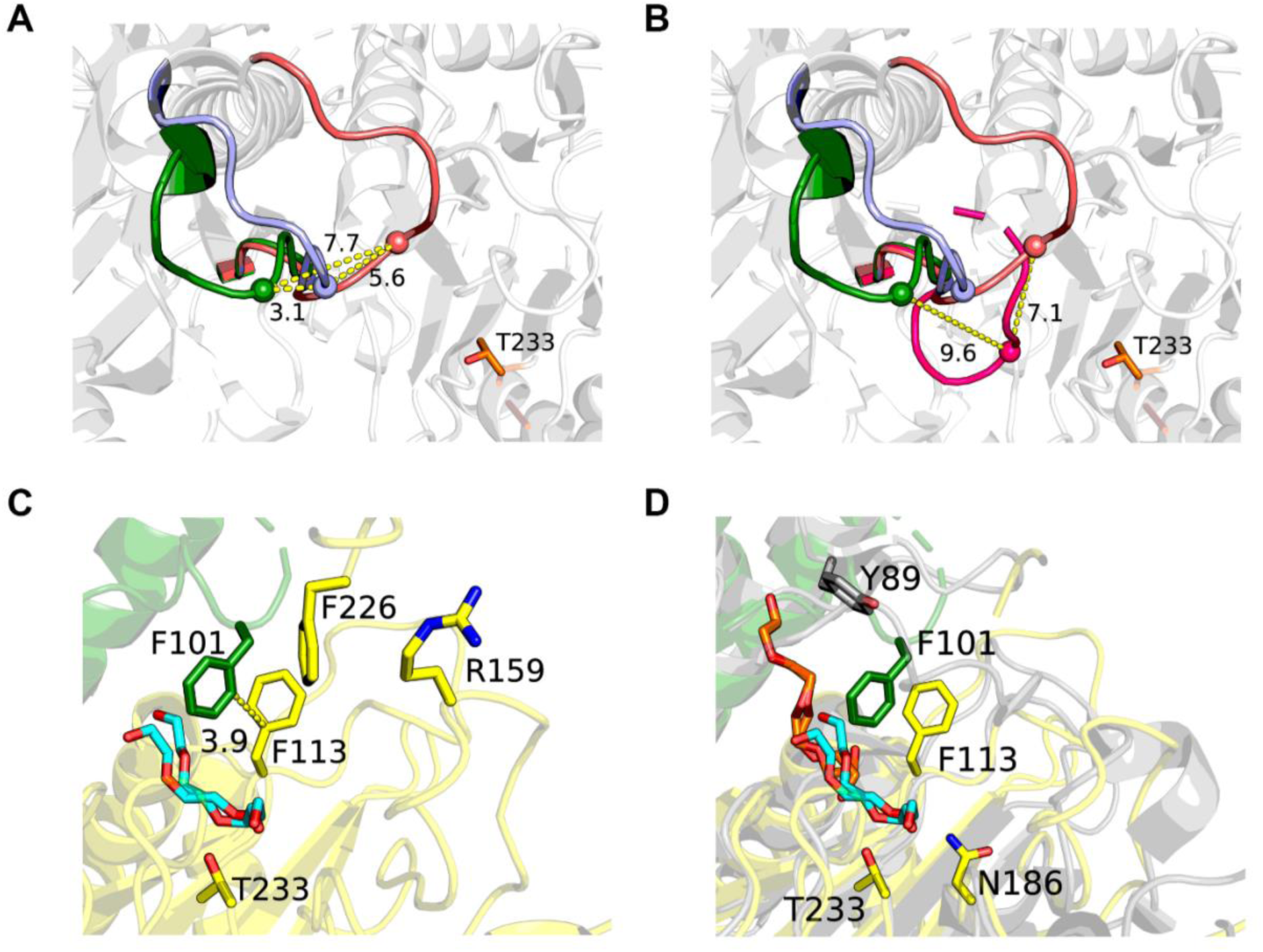
Loop movement in Nyl12. **A.** The loop at the dimer interface of the active sites of Nyl12-CT swings between chain A (A-swing, green), an intermediate position (light blue) and chain D (D-swing, pink). **B.** Comparison between the loop at the dimer interface in Nyl12-CT and Nyl12-RT (hot pink) shows that the latter is bent inward toward the active site, in a “closed” orientation. Distances are represented as dashed lines, measured at the Ca of Phe101 represented as a sphere and expressed in Å. **C.** Residue Phe113 stabilizes the loop in the “closed” conformation in Nyl12-RT, bound to a PEG molecule (cyan). **D.** Comparison of Nyl12-RT and Nyl50-1.8 (PDB ID 9DYS, grey) show a different orientation of the loop and of the aromatic residue harbored on it based on the position of the PEG. Nyl50 appears to have the loop in an “open” conformation due to the binding of the PEG molecule (orange) at a higher position of the active site, with respect to the PEG bound in Nyl12-RT (cyan).

Therefore, to better understand the AS-loop motion with respect to the active site, we generated the tetramers for each assembly found in the asymmetric unit (trimer, A/B dimer, C/D dimer, monomer) with their respective symmetry molecules. Interestingly, the motion of the loop is correlated to the ligand bound state. For the active sites containing a HEPES molecule, the loop is oriented toward the solvent (open conformation), with the Cα of Phe101 at a distance of ∼11-13 Å from the piperazine ring of the HEPES molecule. Conversely, for the active site containing PEG and choline, the loop is bent inward toward the active site (closed conformation) with the Cε of the Phe101 phenyl ring at distances of 3.6 Å and 4.2 Å from C4 and C2, respectively, of the bound PEG molecule.

These extensive movements based on the type of ligand suggests that the AS-loop, located at the dimer interface side of the active site access tunnel, may in fact act as a gate and, considering the similarity between PEG and a linear polyamide, play a major role in substrate positioning with Phe101 as the key residue. Residue Phe101 corresponds topologically to Phe134 in NylC, that has been observed to improve the enzyme’s activity with mutation F134W^20^, highlighting the role of the loop. However, while the NylC-F134W variant showed that the increase in steric hindrance improves the activity via a better amide bond positioning with respect to the catalytic threonine^20^, the same loop in Nyl10 does not harbor aromatic residues but shows a similar hydrolytic activity with a valine residue that topologically corresponds to Phe134, Phe101 and Tyr89 in NylC, Nyl12 and Nyl50, respectively. While residue Phe101 may play a role in substrate positioning, the sidechain of Phe113 appears to form stabilizing interaction with the AS-loop in its closed conformation by forming a T-shaped π-interaction with Phe101 (Figure 4C) and, with the loop in the open conformation, interacts with the piperazine ring of the HEPES molecule with a distance of about 4.5-4.9 Å between the rings (Figure 2). Comparison of the Nyl12-CT and Nyl12-RT PEG-bound dimers to the PEG-bound Nyl50 structure shows that the AS-loop in the latter has adopted an open conformation with the sidechain of Tyr89 oriented towards the solvent. The PEG molecule in the Nyl50 structure is bound in the higher portion of the active site’s access tunnel at the dimer interface (Figure 4D) with the closest atom distant 3.6 Å from the Oγ of catalytic threonine Thr227 and not in an optimal positioning with respect to the oxyanion hole (residues Phe113, Asn180, Thr227) to trigger the downward motion, potentially explaining why the AS-loop adopts an open conformation in this case. A similar loop motion involved in substrate binding was reported for the 6-aminohexanoate dimer hydrolase Hyb-24, in which a mutation stabilizing the loop in the open form improved the catalytic activity of about 10-fold, and alteration of a loop at the active site of the urethanase UMG-SP-1 resulted in a shift in substrate preference.^56,58,59^ Overall, the AS-loop, which is conserved among the nylon oligomer hydrolases, plays a pivotal role in substrate binding and positioning, designating it as an important target for enzyme engineering.

### Ligand-bound Nyl50 structures

To better understand the ligand binding modes and identify the structural determinants responsible for the enzyme’s selectivity, we carried out a series of soaking experiments with a mixture of small soluble linear and cyclic oligomers (PA66 rinsate), obtained after washing the polymer itself with methanol (by-products formed during PA synthesis) and linear monomeric or dimeric products (PA66-L1, -L2). Two Nyl50 structures were determined with non-protein electron density in the active site from soaking experiments with products, both at 3.1 Å resolution. The structure of Nyl50 soaked with PA66-L1 revealed an elongated electron density protruding from the side chain of the catalytic Thr227 residue in both protein chains, suggesting the presence of an acyl-enzyme (Figure S3). We hypothesized that the chemically synthesized products had some carryover longer polyamide molecules that were able to undergo the enzymatic reaction, trapping the enzyme in the acyl-enzyme state. Thus, we modeled and linked a molecule of P66-L1 that is consistent with the electron density (Figure 5A). Superposition of this structure with the previously reported PEG-bound Nyl50-2.2 and 1.8 Å structures (PDB ID 9CXR and 9DYS, respectively) showed that the acylate molecule superimposes with the PEG molecule bound at the dimer interface of the tunnel-like active site in Nyl50-1.8 (Figure 5B). No major changes of the residues in the active site were observed. We also carried out a soaking experiment of a Nyl50 crystal with the commercially available substrate N-(4-nitrophenyl) butanamide (N4NB), which has been used as a control for nylon hydrolytic activity of NylC.^19^ Three different datasets collected at different incubation times (2, 18, and 40 hours) at 1.7, 1.8, and 2 Å resolution revealed an F_O_-F_C_ density near Thr227 that appeared to grow and become more defined with increasing incubation time (Figure S4). The electron density is asymmetric and consistent with a molecule of butyric acid, one of the N4NB hydrolysis products. The carboxylate group is oriented toward the Thr227 side chain, with an average distance of ∼4.3 Å, interacting with the side chain of Ser68 and with the backbone carbonyl oxygen of Asn41 from the same chain (Figure 5C). Similarly to the acylenzyme structure, the butyrate is located at the dimer interface of the active site tunnel. A co-crystal structure of the Ntn-hydrolase *N*-acylethanolamine acid amidase (NAAA) in complex with myristate, one of the products of acylethanolamine hydrolysis, has been reported.^60^ Myristate is bound with its aliphatic tail buried in the hydrophobic portion of the enzyme, with the carboxylate group pointed toward the catalytic cysteine and counterposed to a solvent-exposed arginine residue, an orientation that resembles the one observed for the acylenzyme and the butyrate in Nyl50 despite the structural differences. The computational stage of the Knowvolution approach used a substrate capped on one side to mimic the continuation of the polymer^20^. In a similar manner, the tail of myristate and butyrate can be considered as the capped end and thus the continuation of the polymer. This observation suggests that this subpocket at the dimer interface of the active site is most likely the access site and binding pocket for the substrate.

**Figure 5.**
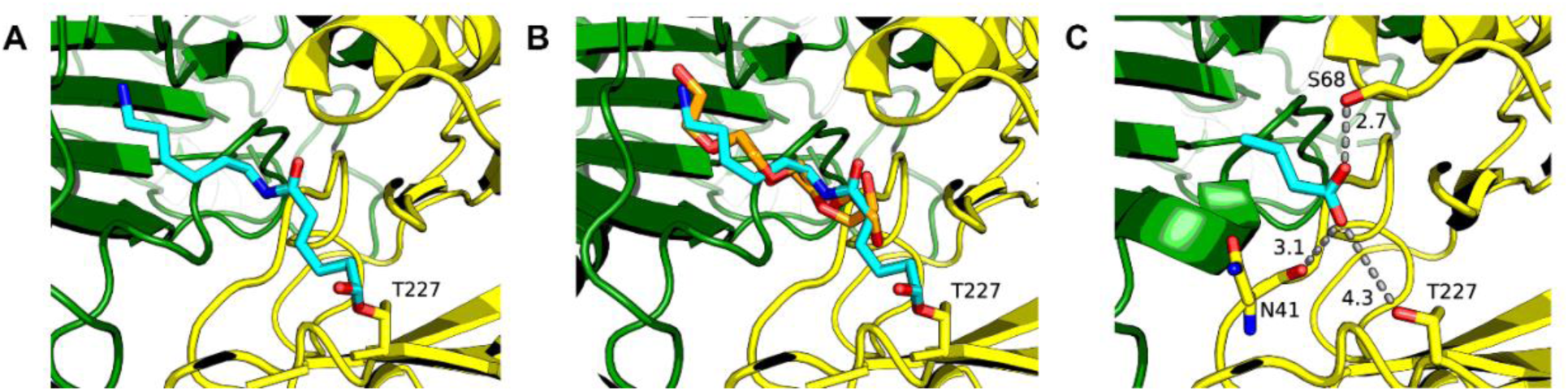
Nyl50 with ligands bound in the active site. **A.** Nyl50 acylenzyme with a PA66 monomer (cyan) covalently bound to Thr227. **B.** Superposition of Nyl50 acylenzyme with PEG-bound (orange) Nyl50-1.8 (PDB ID 9DYS). **C.** Nyl50 bound to butyrate at the A/D dimer interface of the active site, with the substrate carboxylate oriented toward the catalytic Thr227. Color scheme is the same as Figure 1A. Distances (in Å) are shown as dashed lines.

### Charge-dependent substrate selectivity

Based on the orientations of the bound ligands in the nylon hydrolase crystal structures, we envisioned the possibility that the substrate threads inside the active site with its carboxy terminus as the lead rather than the amino group. To test this hypothesis, we adapted the amide hydrolysis colorimetric assay from Bell *et al*.^19^ and used N-(4-nitrophenyl) butanamide (N4NB) and N-(4-aminophenyl) butanamide (N4AB) as substrates. Both N4NB and N4AB are small molecules that contain a butyramide motif similar to the amide bond present in nylon polymers, differing only in the group at the *para* position of the phenyl ring (Figure 6).

**Figure 6.**
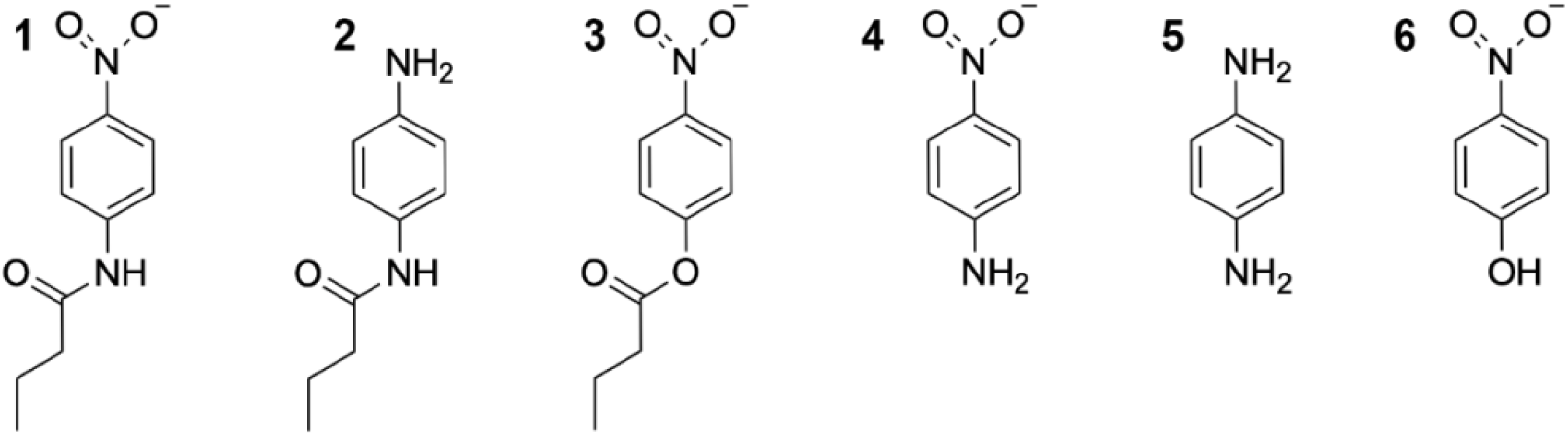
Surrogate substrate molecules used for colorimetric assay: 1. N-(4-nitrophenyl) butanamide (N4NB) **2.** N-(4-aminophenyl) butanamide (N4AB) **3.** N-(4-nitrophenyl) butyrate (N4NBut) **4.** p-nitroaniline (*p*-NA) **5.** p-phenylenediamine (*p*-PD) **6.** p-nitrophenol (*p*-NP).

Hydrolysis of N4NB and N4AB results in the release of *p*-nitronaniline (*p-*NA) and *p*-phenylenediamine (*p-*PD), respectively, which can be followed spectrophotometrically. We observed *p-*NA formation for Nyl10 and Nyl12, and no formation of *p-*PD (Figure S5). However, we did not detect any significant activity for Nyl50 with both substrates, despite monitoring the reaction for 24 hours. Due to the observation of butyrate in the crystal structure of Nyl50 soaked with N4NB, we considered the possibility that the conversion is either very slow or very inefficient under the conditions tested, resulting in insufficient product formation for detection with the colorimetric assay. As a further validation of the binding directionality hypothesis, we tested Nyl10 and Nyl12 against N-(4-nitrophenyl) butyrate (N4NBut) so that it could be used as a non-reactive ligand in crystallization experiments, due to its ester bond. Surprisingly, the activity observed by following the release of N4NBut hydrolysis product *p*-nitrophenol (*p*-NP) was higher compared to the N4NB hydrolysis for both Nyl10 and Nyl12 (Figure S5). Esterolytic activity superior to amide hydrolytic activity was previously reported for Hyb-24^56^, suggesting that these enzymes conserve the preference for ester substrates despite their major structural and sequence differences, thus expanding their possible application to polyesters.

### Computational ligand modeling

We next sought to predict how PA66 binds to Nyl12 using computational modeling. As an alternative to conventional docking, we modeled the enzyme-substrate complex using an approach that combines Boltz-2 ^36^, OpenMM^38^, and PyRosetta^41,42^. Similar to previous work^20^, we capped the N-terminus of PA66-L2 with an N-methylacetamide group to mimic the continuation of the polymer, but the C-terminus was modeled as a carboxylate. Contact restraints were applied during Boltz-2 co-folding to place the desired amide linkage of the substrate near the catalytic residues in the active site. We generated 15 independent enzyme-substrate models, which were then filtered based on Boltz-2 confidence scores and geometric criteria, including selected distances between the substrate and key active site residues (Phe113, Asn186, and Thr233), as well as Bürgi-Dunitz^61,62^ and Flippin-Lodge^63,64^ angles, which define the geometry of nucleophilic attack.

Crystallization additives often bind in the active sites of enzymes, and their placement can provide clues about substrate binding. For example, PEG molecules in the active site of Nyl50 in previously reported X-ray crystal structures were used to help identify the active site tunnel.^15^ Similarly, the HEPES, acetate, and PEG molecules observed in the Nyl12 structures reported in the present work reveal structural determinants that may be involved in substrate binding and expulsion. Thus, as an additional validation step, we superimposed each enzyme-substrate model onto three X-ray crystal structures: Nyl12-CT, and Nyl50 PA66 acylenzyme, and a PEG-bound structure of Nyl50 (PDB ID 9CXR). Eight of the 15 models passed the geometric filters, and out of these, in six models the substrates were generally oriented with the terminal carboxylate located at the portion of the tunnel that is exposed to the solvent and interacting with residue Arg159 via the carboxylate or an amide group. Several models displayed similar conformations on one portion of the substrate (e.g., the carboxylate portion, from the scissile amide bond) while the other portion folded over itself. Two models were deemed the most accurate but only one had the best fit when compared to the tunnel and the PEG molecules in the Nyl50 structures with the only difference being the orientation of the alkyl chains (Figure 7A). We found that i) a substrate amide N in the top Nyl12-PA66 model superimposes onto the choline N to within 1.8 Å (Figure 7B), ii) Arg159 is hydrogen bonded to a substrate’s amide oxygen and superimposes onto Arg159 in Nyl12-CT, and iii) Alkyl chains in the top model superimpose onto PEG in a Nyl50 structure (PDB ID 9CXR) and partially with the acyl chain of the Nyl50 acylenzyme (Figure 7C).

**Figure 7.**
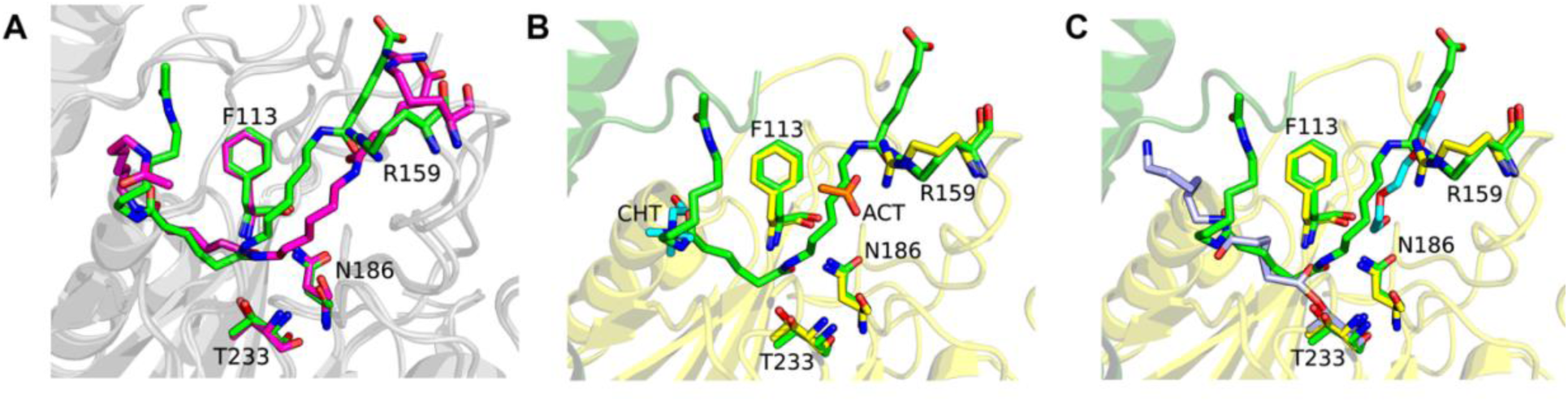
**A.** Comparison of the top two computational models of Nyl12-PA66 (green and magenta) with selected residues and the PA66 substrate shown as sticks. **B.** Superposition of the best-fitting model (green sticks) onto the Nyl12-CT (cartoon and yellow sticks) with choline (CHT, cyan) and acetate (ACT, orange) bound near the active site. An amide bond of the substrate overlaps with the N atom of choline. Arg159 in Nyl12-CT is hydrogen bonded to the acetate molecule. **C.** Superposition of the model onto the X-ray structure of Nyl50 acylenzyme (light blue), Nyl50 with PEG (cyan) bound near the active site (PDB ID 9CXR), and Nyl12 (cartoon, green and yellow sticks). Arg159 has a similar orientation in the crystal structure and the model, in which it interacts with an acetate molecule and an amide bond of the PA66 substrate, respectively.

Overall, the top models suggest a possible substrate binding pose (Figure S6) that is validated by comparison with non-substrate bound X-ray crystal structures. The comparison informs on the quality of the models, which appear to be accurate up to the amide bond that superimposes with the choline molecule, after which the positioning becomes less consistent, also considering the orientation of the PEG molecules bound at the dimer interface of Nyl50 and Nyl12 structures. The two different orientations of Arg159 based on the position of the carboxylate group of the substrate suggest that this residue may play a role in stabilizing the substrate and guiding the leaving group after the hydrolysis. The observed consensus of the orientation of the terminal carboxylate in the models supports the hypothesis that the enzyme threads the substrate into the active site with carboxylate terminus of the substrate as the lead.

### Oligomeric assembly of nylon hydrolases

Previous studies have reported that the quaternary structures of nylon hydrolases are a mixture of oligomeric states in solution^15,16^, with most proposed to be in equilibrium between the dimeric and tetrameric forms.^16^ Although the functional unit is the dimeric form, NylC from different species was observed to exist in solution as a tetramer, in an equilibrium of dimers and tetramers, and as a mixture of dimer, trimer and tetramer for wild-type NylC_A_, NylC_K_ and NylC_p2_, respectively.^16^ It is not uncommon for members of the Ntn-hydrolase superfamily to assemble in different oligomeric states. For example, the human L-asparaginase-like protein 1 (ASRGL1) is a trimer in solution.^65^ Oligomerization is a prerequisite for autocatalytic cleavage and is also involved in maintaining protein stability.^16,65^ Previously, we reported Nyl50 as a mixture of dimers and tetramers in solution, depending on concentration.^15^ Because we observed a discrepancy in the oligomeric state among size-exclusion chromatography, DLS, and AUC, we used a series of biophysical methods to determine the oligomeric state for Nyl10, Nyl12 and Nyl50 in solution. The chromatography profiles were obtained from analytical SEC, and the MW was calculated after calibrating the column to obtain a relation between MW and elution volume. For all three nylon hydrolases, the elution occurred in a single, sharp peak, at an elution volume between 1.42 and 1.46 mL, consistent with the MW of a dimer (Figure 8A). Next, we carried out a DLS analysis to assess the polydispersity of the protein samples prior to the crystallization experiments. Again, all three Nyl10, Nyl12 and Nyl50 samples, showed low polydispersity (<15%) and radius of about 3.6-3.9 nm, with a calculated MW between 60 and 80 kDa, which corresponds approximatively to the size of a dimer (Figure S7).

**Figure 8.**
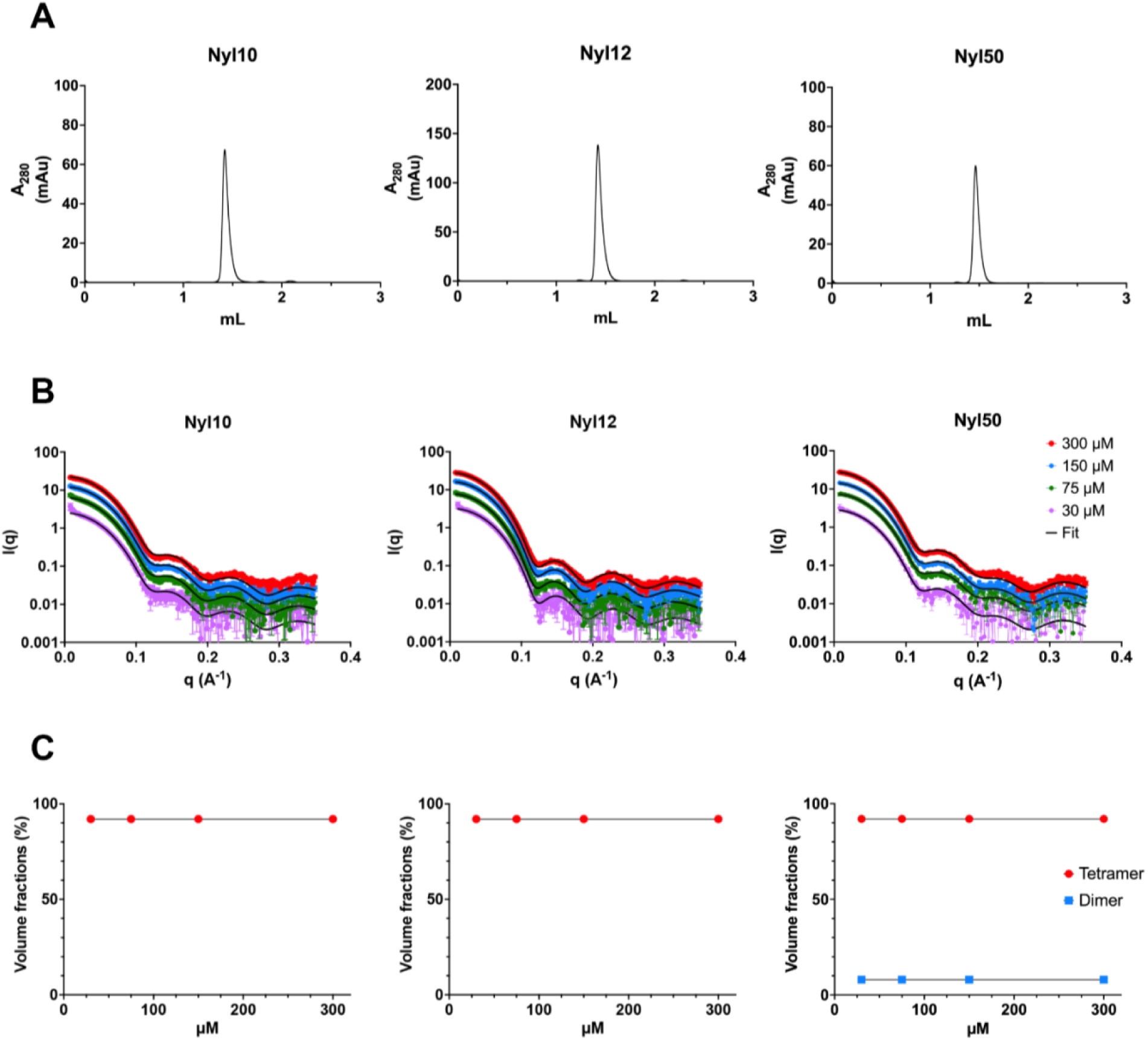
**A.** Size-exclusion chromatography profiles of the studied nylon hydrolases. **B.** Small Angle X-ray Scattering (SAXS) profiles of nylon hydrolases at different concentrations, superimposed to the theoretical fit (black line) for a tetramer, showing the same profile at different concentration and a good agreement with the theoretical fit. **C.** Oligomeric distribution calculated by the software OLIGOMER at different concentrations shows that the tetrameric assembly is the predominant assembly independent of concentration.

To investigate if the oligomeric state is concentration-dependent, we carried out a SAXS analysis. Using four concentrations between 30 and 300 μM (1 and 10 mg/mL), we encompassed the working concentrations of purification, DLS, and crystallization. Average MW was calculated directly from experimental data using the forward scattering intensity, *I*(0) (Table S3), and subsequently with indirect methods using both experimental data and high-resolution crystallographic structures. The latter approach allowed for the calculation of the theoretical scattering profile and radius of gyration (R_g_) based on the dimeric and tetrameric crystal structures using the FoXS webserver^47,48^, and the determination of volume fractions of the components (form factors) from the SAXS data using the OLIGOMER^49^ program. Scattering profiles were similar for all three studied nylon hydrolases, suggesting a tetramer in solution across all measured concentrations (Table S3). Furthermore, a fit of the tetrameric crystal structure agrees with the scattering data at each concentration point (Figure 8B). Oligomeric distribution analysis for Nyl50 calculated the presence of tetrameric and dimeric species with an average volume fraction of 92% for the tetramer, while for Nyl10 and Nyl12 the oligomeric distribution was consistent with the sole presence of a tetrameric species across all analyzed concentrations (Figure 8C).

Next, to understand if a lower-order quaternary structure was being masked by the scattering profile of the tetrameric species, we carried out AUC sedimentation velocity analysis at 30, 15, and 3 μM (1, 0.5 and 0.1 mg/mL). For all three enzymes, AUC revealed a major, sharp peak with an average sedimentation coefficient (*S*) of 6.5 for Nyl10 and 7.1 for Nyl12 and Nyl50, congruent with the MW of a tetramer (Figure 9A), accounting for ∼98% of the sample at all concentrations for Nyl10, and more than 90% of the sample for Nyl12 and Nyl50 at 30 and 15 μM concentrations. A small bump was observed at a coefficient of ∼10 *S*, corresponding to a calculated MW of 230 kDa, that increases in intensity as the concentration diminishes, which we ascribe to aggregation. Interestingly, a small bump at ∼3 *S*, corresponding to the size of a monomer (35 kDa), was observed at a concentration of 3 μM for Nyl12 and accounts for 1% of the population (Figure S8-S10).

**Figure 9.**
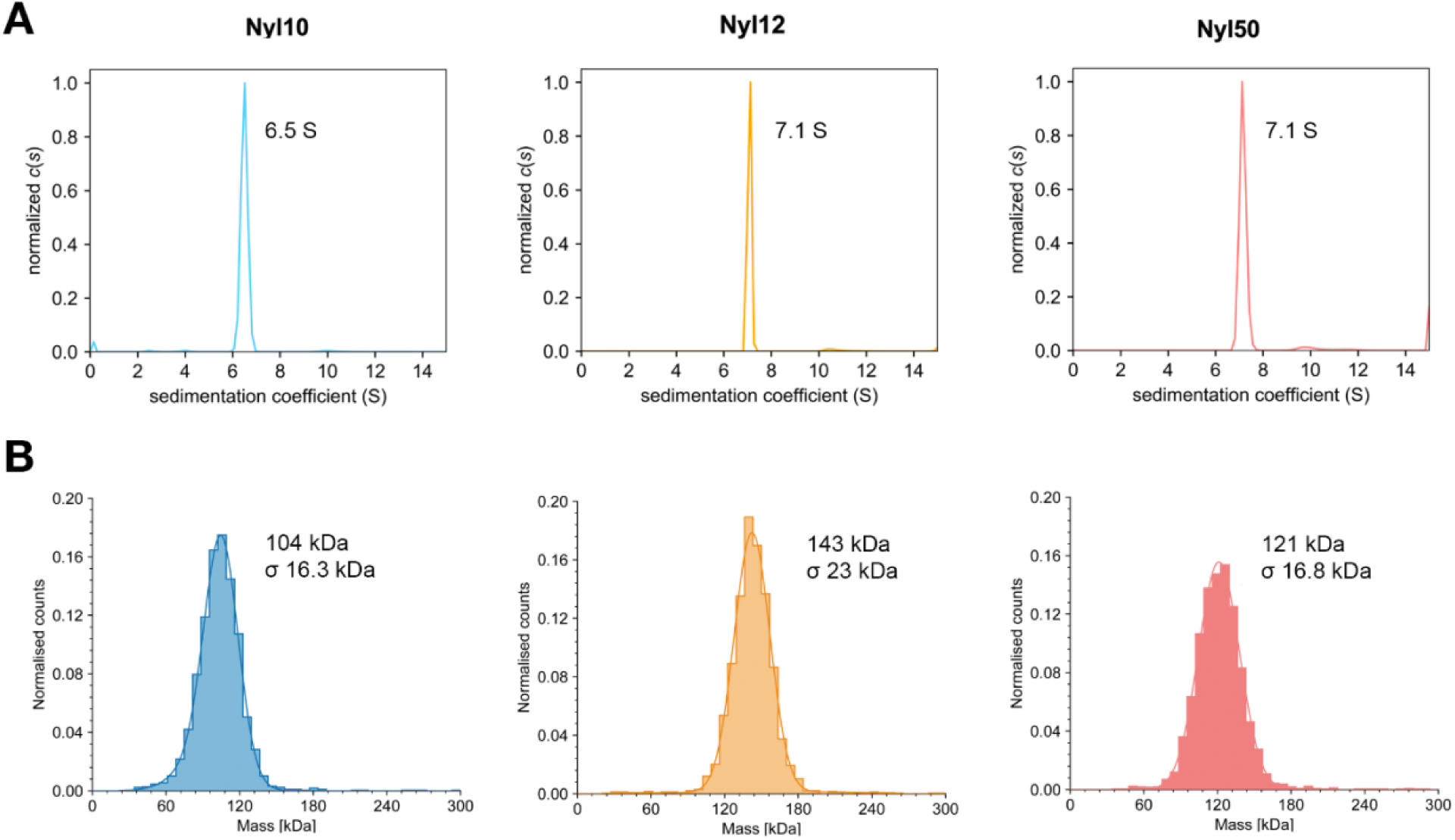
Assessment of oligomeric assembly. **A.** Normalized sedimentation velocity profiles by analytical ultracentrifugation (AUC) of Nyl10, Nyl12 and Nyl50 showing a single peak, that is consistent with the size of a tetramer. **B.** Mass photometry analysis showing a single peak, consistent with a tetrameric form.

We then investigated if the oligomeric distribution would also be retained at a very low concentration. We performed mass photometry (MP) analysis using a single, low enzyme concentration of ∼20 nM. MP analysis is restricted to low concentrations (<100 nM) due to instrument limitations and limited to the detection of macromolecules larger than 30 kDa, which would make the detection of monomeric states difficult. However, we observed only higher order oligomers, with a single peak corresponding to the tetrameric form for all three nylon hydrolases (Figure 9B).

The discrepancy between the MW observed in SEC and DLS analyses and the SAXS, AUC and MP analyses, can be explained by the limitations of the first two methods. To estimate the protein’s MW, both SEC and DLS rely on hydrodynamic radius (R_h_), which is dictated not only by the mass of the protein but by the shape as well. Conversely, SAXS, AUC, and MP are matrix-free, direct measurements methods, capable of measuring the physical properties of the proteins and thus less prone to inaccuracies caused by non-globular shapes and column interactions. Therefore, we suspect that these two methods cannot discern between the dimeric and tetrameric form due to an R_h_ that is similar between the species. We note that it is not uncommon to observe differences in oligomeric states based on the method used.^66^

These findings suggest that the tetrameric quaternary structure is the predominant assembly in solution for all three nylon hydrolases, independent of concentration. Knowing the correct quaternary structure helps avoid pitfalls when identifying and analyzing the active site of a multimeric protein, particularly if no ligand-bound crystal structures are present. Thus, combining this information with the crystal structure analysis and the residues that interact with the ligands in the active site, we hypothesize that the tetramer is formed and conserved due to the participation of residues from a third subunit in the formation of the active site.

### Comparative catalytic efficiency of nylon hydrolases toward PA66

We then compared the catalytic activities of these three enzymes by determining their Michaelis-Menten kinetic parameters using the conventional method to maintain consistency with previous studies.^20,53^ Enzymatic reactions were conducted under optimal conditions (75 °C, 200 mM Tris buffer, pH 8) for 2 hours, as established in our previous work.^54^ Hydrolysis products were analyzed via I.DOT/OPSI-MS.

Due to PA66 substrate specificity for Nyl10 and Nyl50, kinetic parameters were measured using PA66, marking the first report of such parameters determined on a substrate other than PA6. The data are summarized in Table 1 and Figure S11 A-C. Overall, Nyl50 and Nyl10 exhibited comparable kinetic parameters, while Nyl12 demonstrated significantly enhanced catalytic performance. Specifically, Nyl12 showed a V_max_ approximately 10-fold greater (0.318 μM·s⁻¹) and a k_cat_ about 5.5-fold higher (1.72 s⁻¹) than those of the homologous enzymes (Figure S11). Ultimately, despite having three times less affinity for PA66 compared to Nyl50, Nyl12 is about 2- and 5-fold more efficient than Nyl50 and Nyl10, respectively. These findings align with our recent observations where Nyl12 achieved the higher degree of depolymerization of PA66 (∼2%), surpassing its homologs (∼0.5%).^54^

**Table 1.** Kinetic parameters of Nyl50, Nyl10, and Nyl12 with PA66 and comparison with NylC_TS_ and NylC_TS_^P27Q/F301L^.

| Substrate | Enzyme | $V_{\max}$ ($\mu\text{M}\cdot\text{s}^{-1}$) | $k_{\text{cat}}$ ($\text{s}^{-1}$) | $K_{\text{M}}$ ($\text{g}\cdot\text{L}^{-1}$) | $k_{\text{cat}}/K_{\text{M}}$ ($\text{L}\cdot\text{s}^{-1}\cdot\text{g}^{-1}$) | Source |
| --- | --- | --- | --- | --- | --- | --- |
| PA66 | Nyl50 | 0.040 | 0.32 | 82.2 | 0.0039 | This study |
|  | Nyl10 | 0.035 | 0.29 | 201.9 | 0.0015 | This study |
|  | Nyl12 | 0.318 | 1.72 | 231.1 | 0.0074 | This study |
| PA6 | NylC<sub>TS</sub><sup>P27Q/F301L</sup> | 0.053 | 1.06 | 114.8 | 0.0092 | Puetz et al.<sup>53</sup> |
|  | NylC<sub>TS</sub> | 0.028 | 0.57 | 109 | 0.0052 | Puetz et al.<sup>53</sup> |

Moreover, the kinetic parameters of Nyl12 are comparable to previously reported values for PA6, with Nyl12 performing slightly better than NylC_TS_ and the NylC_TS_^P27Q/F301L^ mutant (Table1). Direct comparisons are complicated by differences in substrate type (PA66 vs. PA6) and form (powder vs. film).

The wild-type Nyl12 enzyme appears to outperform (higher titer) its engineered homologs.^54^ Together with the structural insights presented above, this makes it an excellent candidate for future enzyme engineering campaigns.

### Conclusions

This study presents the first crystal structures of two product- and substrate-selective nylon hydrolases, named Nyl10 and Nyl12, and two ligand-bound structures of the Nyl50 nylon hydrolase. Structural analysis revealed that a flexible loop proximal to the active site, present in all the analyzed nylon hydrolases, adopts “inward” and “outward” conformations depending on substrate binding. This motion suggests an induced-fit substrate binding mechanism. The role of this loop in substrate stabilization is supported by analogy to an engineered variant of NylB, the 6-aminohexanoic linear dimer hydrolase Hyb-24, and the urethanase UMG-SP-1. Mutations targeting residues on the corresponding loop of Hyb-24 and UMG-SP-1 enhanced the amidase activity.^20,56,59^

Furthermore, we gained insight into polyamide binding directionality into the active site tunnel using short soluble substrates and computational modeling. Namely, the polyamide substrate would access the active site tunnel at the dimer interface and with the carboxylic group leading.

Comprehensive solution characterization established that all three nylon hydrolases, including Nyl50, adopt a tetrameric quaternary structure independent of protein concentration. The combination of the structural data and the information on the oligomeric assembly strongly indicates that the active site is formed by the participation of three protomers, providing a valuable insight that will help protein engineering strategies aimed to enhance amidase activity or shift substrate preference, whether by classic or computational approaches.

Based on crystallographic evidence, we hypothesize that the polyamide substrate is hydrolyzed in a processive manner, via the following reaction model: considering the A/D dimer, the polymer strand (i) enters via monomer A, threading all the way in the active site and fully occupying the tunnel, (ii) the AS-loop moves down and stabilizes the substrate, (iii) the polymer is cleaved and the product leaves via the monomer D, possibly with Arg159 (Nyl12 numbering) acting as a “hook” on the carboxylate terminal group. Cleavage of the amide bond exposes a carboxylate group to the hydrophobic portion of the active site, (iv) the AS-loop lifts and the polymer slides forward until the correct geometry for cleavage is reached and the reaction is repeated.

Kinetic assays showed that the catalytic efficiency of Nyl12 with PA66 is comparable to the engineered NylC variant, NylC ^P27Q/F301L^ with its best substrate PA6, while it is currently the enzyme producing the highest titer.^54^ Thus, Nyl12 is a promising candidate for protein engineering for efficient nylon hydrolysis. Activity assays showed that both Nyl10 and Nyl12 are also capable of hydrolyzing ester bonds, as it has been observed for Hyb-24 despite the very different structure, introducing the possibility of the application of nylon hydrolases and nylon enzymatic recycling protocols to polyesters.

In conclusion, we determined crystal structures of two novel nylon hydrolases, elucidated the oligomerization state of relevant NylC homologs in solution, assessed the kinetic parameters of these enzymes with PA66 substrate, investigated substrate binding directionality, and combined the evidence to propose a model describing the enzymatic reaction steps. Overall, these findings provide insights into the mechanism of nylon hydrolases, further understanding of the role of the structural determinants and quaternary structure, provide starting points for engineering catalytic activity and specificity in nylon hydrolases, and identify Nyl12 as a versatile candidate for future protein engineering campaigns.

## Acknowledgements

We thank Dr. S. Khan and Dr W.C. Leite for the support with the SAXS and AUC experiments and analysis, and I. Mathews for support during data collection at the Stanford Synchrotron Radiation Lightsource. Crystallization at the National Crystallization Center at the Hauptman-Woodward Institute was supported through NIH grant R24GM141256.

## Author contribution

N.C.: conceptualization, enzyme expression, colorimetric assays, crystallization, structural characterization and biophysical analysis, analysis of computational models; C.B.: Enzyme kinetics and analysis; J.M.P.: Computational ligand modeling and analysis; D.L.C: Analytical characterization; J.F.C: Analytical methods development and analysis; supervision of analytical research J.K.M.: funding acquisition, project management; F.M.: conceptualization, structural analysis. All authors contributed to writing and editing the manuscript.

