## Supplementary Information for "Structural and oligomeric characterization of substrate- and product-selective nylon hydrolases"

| **Table S1: Data collection and refinement statistics** | | | | | |
| --- | --- | --- | --- | --- | --- |
|  | **Nyl10** | **Nyl12-RT** | **Nyl12-CT** | **Nyl50-acylenzyme** | **Nyl50-N4NB** |
| **Matthews Coefficient (solvent content, %)** | 2.13 (42) | 2.39 (48.5) | 2.06 (40.3) | 2.18 (43.6) | 2.18 (43.6) |
| **Data collection** |  |  |  |  |  |
| Beamline | SSRL BL 12-1 | In-house | SSRL BL 12-1 | In-house | In-house |
| Wavelength (Å) | 0.98 | 1.54 | 0.98 | 1.54 | 1.54 |
| Space group | C 2 | P 2 2_1_ 2_1_ | P 2_1_ 2_1_ 2_1_ | P 2 2_1_ 2_1_ | P 2 2_1_ 2_1_ |
| Unit cell dimensions |  |  |  |  |  |
| a, b, c (Å) | 93.13, 120.19, 148.30 | 77.59, 79.43, 108.77 | 119.62, 135.41, 142.88 | 53.81, 96.45, 104.66 | 53.88, 96.48, 105.46 |
| α, β_,_ γ (°) | 90, 93, 90 | 90, 90 ,90 | 90, 90, 90 | 90, 90, 90 | 90, 90, 90 |
| Resolution range (Å)^*^ | 74.05 - 1.30 (1.32-1.30) | 29.64 - 1.90 (1.94-1.90) | 98.28 - 1.75 (1.78 - 1.75) | 27.39 - 3.10 (3.31 - 3.10) | 28.41 - 2.0 (2.05-2.0) |
| Total reflections^*^ | 1357612 (67094) | 314936 (14308) | 1486470 (71568) | 98455 (17990) | 236423 (17590) |
| Unique reflections^*^ | 351281 (17449) | 53622 (3584) | 217968 (11205) | 10389 (1842) | 36372 (2573) |
| <I/σ>^*^ | 10.7 (1.4) | 13.3 (2.1) | 8.2 (1.5) | 9.9 (4.5) | 8.7 (1.5) |
| CC_(1/2)_ ^*^ | 0.99 (0.44) | 0.99 (0.72) | 0.99 (0.65) | 0.98 (0.90) | 0.97 (0.47) |
| Completeness (%)^*^ | 98.8 (99.2) | 99.9 (99.9) | 93.7 (98.0) | 99.8 (100.0) | 96.1 (94.4) |
| Wilson B-factor (Å^2^) ^*^ | 14.5 | 23.5 | 16.6 | 13.1 | 22.2 |
| R_merge_ | 0.051 (1.165) | 0.063 (0.717) | 0.134 (1.177) | 0.265 (0.587) | 0.172 (1.76) |
| **Refinement** |  |  |  |  |  |
| R_work_/ R_free_^#^ | 0.14/0.17 | 0.15/0.20 | 0.16/0.20 | 0.14/0.22 | 0.17/0.22 |
| Number of non-H atoms in AU^†^ | 12427 | 4873 | 19711 | 4007 | 4400 |
| protein | 11664 | 4657 | 18757 | 4373 | 4317 |
| ligands | 45 | 34 | 119 | 34 | 12 |
| solvent | 788 | 182 | 835 | 0 | 71 |
| Average B-factor (Å^2^) | 15.4 | 31.1 | 22.42 | 8 | 31.04 |
| RMSD |  |  |  |  |  |
| Bond lengths (Å) | 0.012 | 0.016 | 0.013 | 0.013 | 0.015 |
| Bond angles (Å) | 1.71 | 2.29 | 2.07 | 1.93 | 2.34 |
| Ramachandran plot |  |  |  |  |  |
| % favored, outliers | 97.71, 0.0 | 97.0, 0.0 | 97.8, 0.04 | 97.14, 0.0 | 97.11, 0.0 |
| Clashscore | 2.19 | 4.46 | 3.14 | 3.96 | 2.18 |
| PDB ID | 9YYS | 9Z0A | 9Z0B | 9Z0Z | 9Z11 |
| AU is asymmetric unit  ^*^ Values in parentheses correspond to the highest resolution shell.  ^#^ R_free_ is calculated as R_work_ using 5% of all reflections randomly chosen, which were excluded from structure refinement.  ^†^The asymmetric unit contained 6, 2, 8, and 2 protein molecules for Nyl10, Nyl12-RTNyl12-CT, Nyl50-acylenzyme and Nyl50-N4NB, respectively. | | | |  |  |

| **Table S2: List of residues forming the active site** | | | |
| --- | --- | --- | --- |
| **Monomer A** | **Nyl50** | **Nyl10** | **Nyl12** |
|  | 49 LEU | 50 ALA | 57 GLN |
|  | 57 PRO | 58 THR | 65 ALA |
|  | 58 SER | 59 ALA | 66 TYR |
|  | 59 VAL | 60 VAL | 67 LEU |
|  | 60 THR | 61 ASP | 68 ASP |
|  | 88 GLY | 89 GLY | 100 GLN |
|  | 89 TYR | 90 VAL | 101 PHE |
|  | 96 ILE | 97 VAL | 104 MET |
|  | 97 PRO | 98 PRO | 105 PRO |
|  | 98 LEU | 99 ILE | 106 LEU |
| **Monomer D** | 41 ASN | 42 ALA | 50 SER |
|  | 42 ALA | 43 PRO | 51 VAL |
|  | 43 ALA | 44 ALA | 52 GLY |
|  | 68 SER | 69 SER | 76 GLY |
|  | 103 VAL | 104 ALA | 111 VAL |
|  | 104 ILE | 105 LEU | 112 ILE |
|  | 105 TYR | 106 TYR | 113 PHE |
|  | 108 SER | 109 ALA | 116 THR |
|  | 148 LYS | 149 ASN | 156 LYS |
|  | 151 GLY | 152 GLY | 161 GLY |
|  | 152 VAL | 153 VAL | 162 PHE |
|  | 180 ASN | 180 ASN | 186 ASN |
|  | 227 THR | 211 THR | 233 THR |
|  | 265 VAL | 249 VAL | 271 LEU |
|  | 267 GLY | 251 GLY | 273 GLY |

| **Table S3. Summary of Small-angle X-ray scattering results** | | |  |
| --- | --- | --- | --- |
| **Sample** | Conc (mg/ml) | R_g_ (Å) | d_max_ (Å) |
| **Nyl10** |  |  |  |
|  | 1 | 29.78 ± 0.1 | 90 |
|  | 2.5 | 29.29 ± 0.07 | 90 |
|  | 5 | 28.71 ± 0.04 | 95 |
|  | 10 | 29.08 ± 0.01 | 85 |
| **Nyl12** |  |  |  |
|  | 1 | 30.67 ± 0.07 | 92 |
|  | 2.5 | 30.5 ± 0.03 | 90 |
|  | 5 | 30.41± 0.02 | 90 |
|  | 10 | 30.31 ± 0.02 | 95 |
| **Nyl50** |  |  |  |
|  | 1 | 30.48 ± 0.08 | 95 |
|  | 2.5 | 29.71 ± 0.04 | 90 |
|  | 5 | 29.65 ± 0.03 | 94 |
|  | 10 | 29.05 ± 0.02 | 95 |
| **Structural parameters^a^** | **Nyl10** | **Nyl12** | **Nyl50** |
| Theoretical R_g_ (FoXS, tetramer) | 27.71 | 28.56 | 28.77 |
| Fit χ^2^ (FoXS) | 2.19 | 3.01 | 1.97 |
| **Guinier analysis** |  |  |  |
| *I*(0) | 6.45 ± 0.02 | 8.19 ± 0.02 | 7.55 ± 0.02 |
| R_g_ (Å) | 29.95 ± 0.11 | 31.49 ± 0.1 | 30.53 ± 0.1 |
| *q*-range | 0.018 - 0.04 | 0.015 - 0.04 | 0.015 - 0.04 |
| ***P(r)* analysis** |  |  |  |
| *I*(0) | 6.39 ± 0.01 | 8.05 ± 0.01 | 7.46 ± 0.01 |
| R_g_ (Å) | 29.29 ± 0.07 | 30.5 ± 0.03 | 29.71 ± 0.04 |
| d_max_ (Å) | 90 | 90 | 90 |
| *q*-range (Å^-1^) | 0.018 - 0.4 | 0.015 - 0.26 | 0.014 - 0.3 |
| χ^2^ (GNOM) | 1.15 | 1.51 | 1.39 |
| **Molecular weight analysis** |  |  |  |
| Calculated weight (kDa) | 28.9 | 35.1 | 31.4 |
| Experimental Vc (kDa) | 114.9 | 139.6 | 126.8 |
| *q*_max_ cutoff (Å^-1^) | 0.4 | 0.4 | 0.4 |
| **OLIGOMER fitting** |  |  |  |
| Multimers used | Dimer, tetramer | Dimer, tetramer | Dimer, tetramer |
| Oligomer R_g_ (dimer, tetramer) | 23.97, 28.85 | 26.25, 29.53 | 25.06, 29.95 |
| *q*-range for fitting (Å^-1^) | 0.01 – 0.4 | 0.01 – 0.4 | 0.01 – 0.4 |
| χ^2^ | 8.70 | 9.18 | 3.04 |
| ^a^parameters reported for sample concentration at 2.5 mg/ml | | | |

**Figure S1**. **A**. Multiple sequence alignment of Nylon hydrolases colored by identity. **B**. Graphical representation of sequence identity among the studied nylon hydrolases.

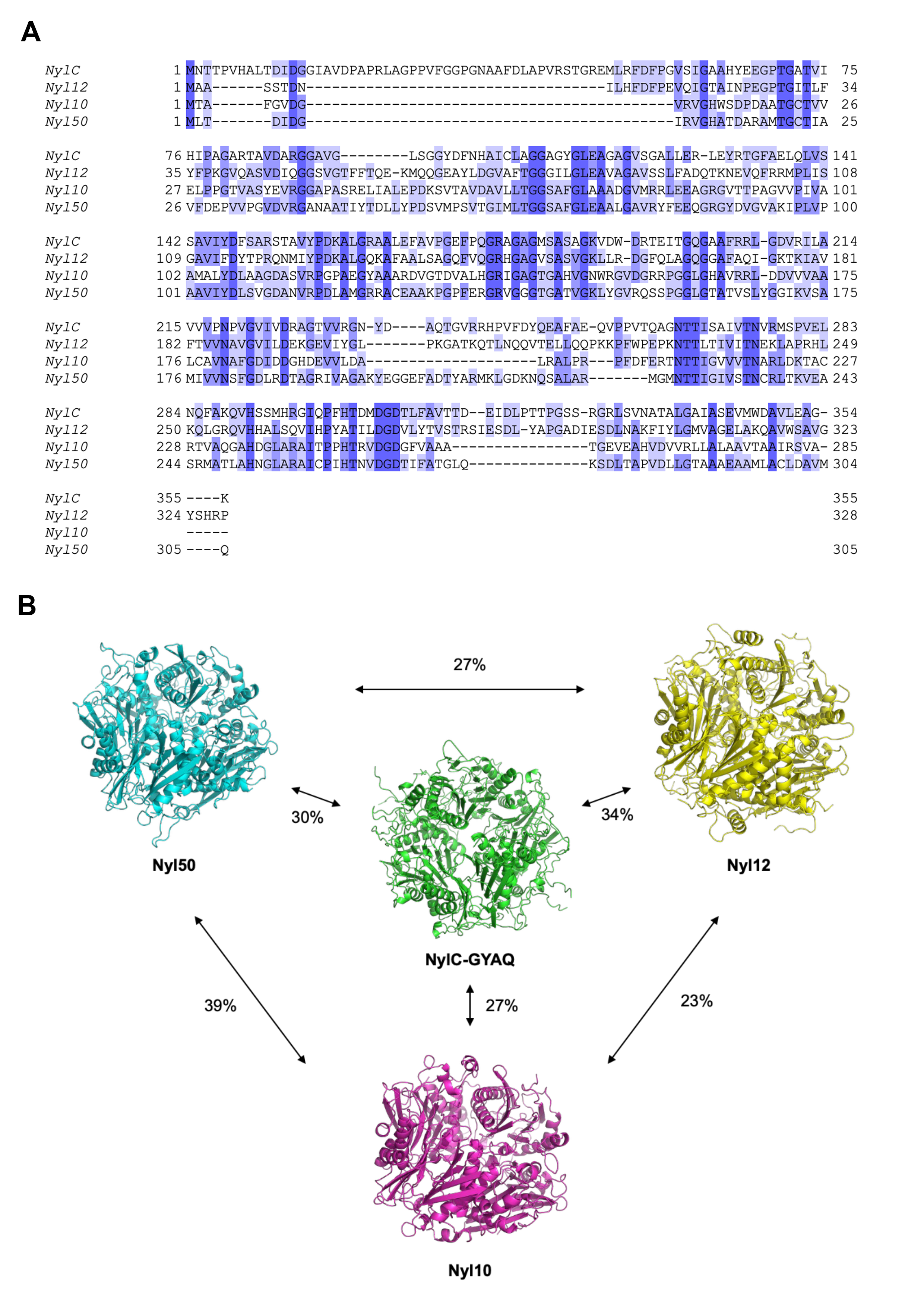

**Figure S2**. **Crystallization experiments of Nyl12. A.** Crystals obtained by under oil micro batch crystallization experiments from which Nyl12-RT datasets were collected. **B.** Crystals obtained by liquid-gel diffusion in a 1.5 mL Eppendorf tube from which Nyl12-CT datasets were collected. Notably, the different experiment setup influenced the morphology of the crystal.
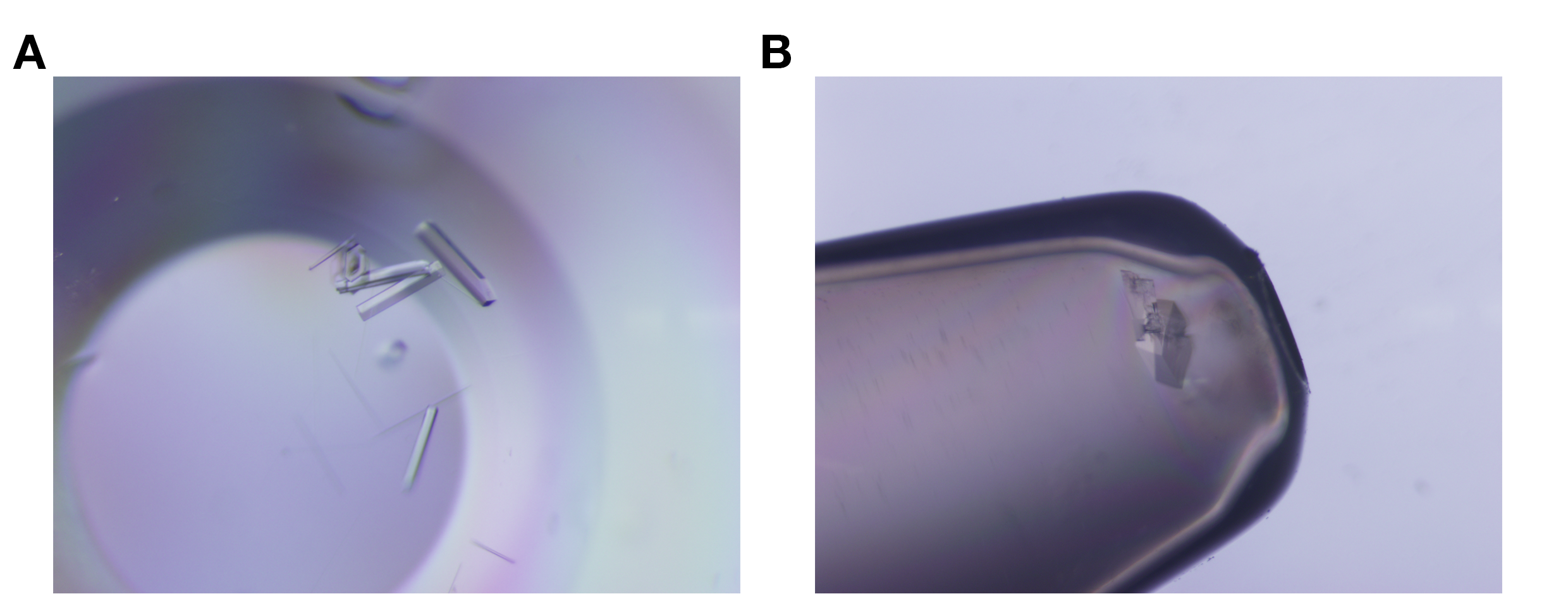

**Figure S3**. **A**. Continuous electron density showing extruding from catalytic Thr227 in Nyl50 structure at 3.1 Å resolution. 2F_O_-F_C_ map is contoured at 1.0 σ and carved at 2.0 Å. **B**. Coot view of electron density protruding from Thr227, 2F_O_-F_C_ map is contoured at 1.0and F_O_-F_C_ contoured at 3.0 σ.

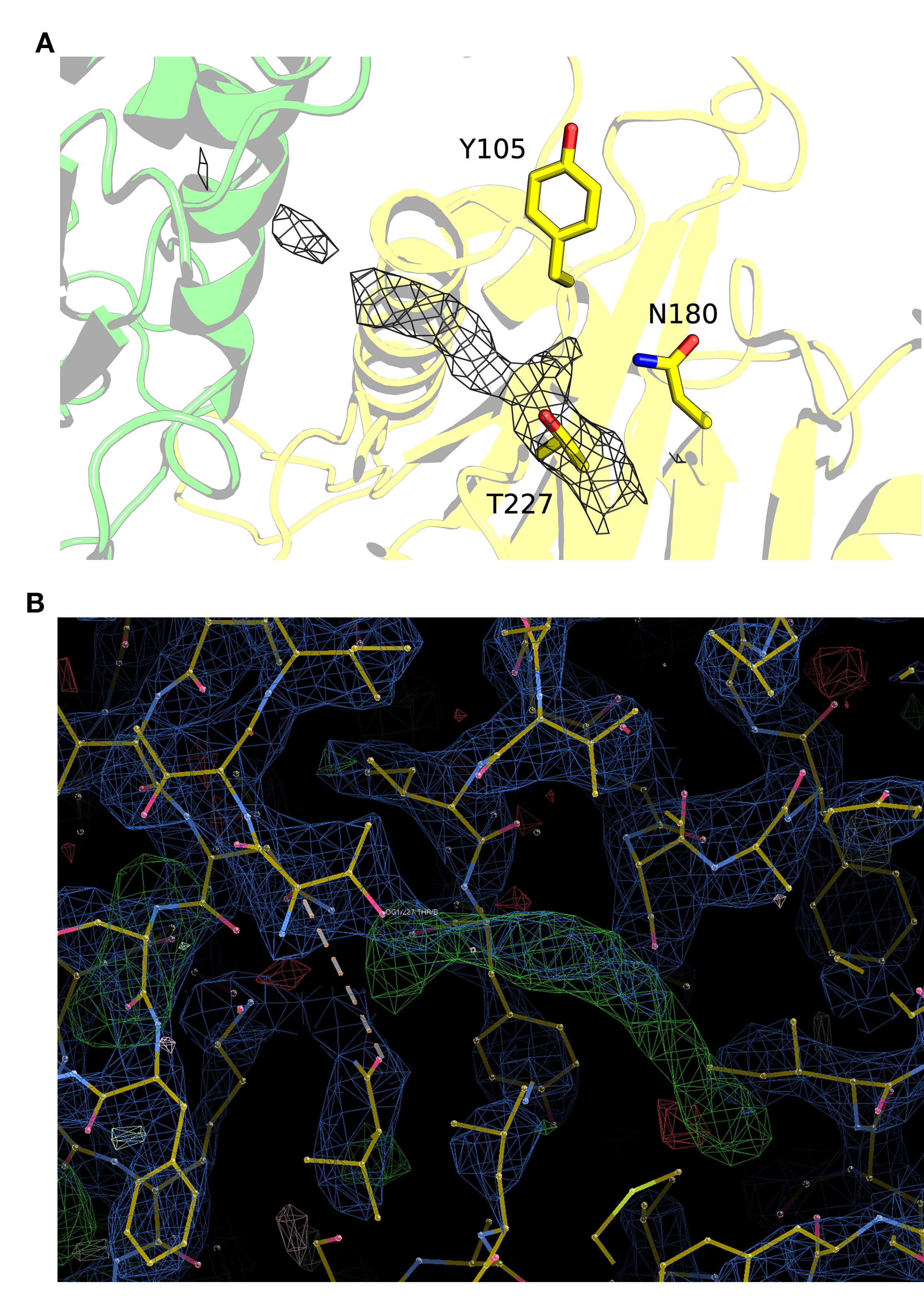

**Figure S4**. **A-C**. Electron density of Nyl50 crystal soaked with N4NB after 2 hours, overnight and 40 hours at 1.85, 2.0 and 2.3 Å resolution, respectively.

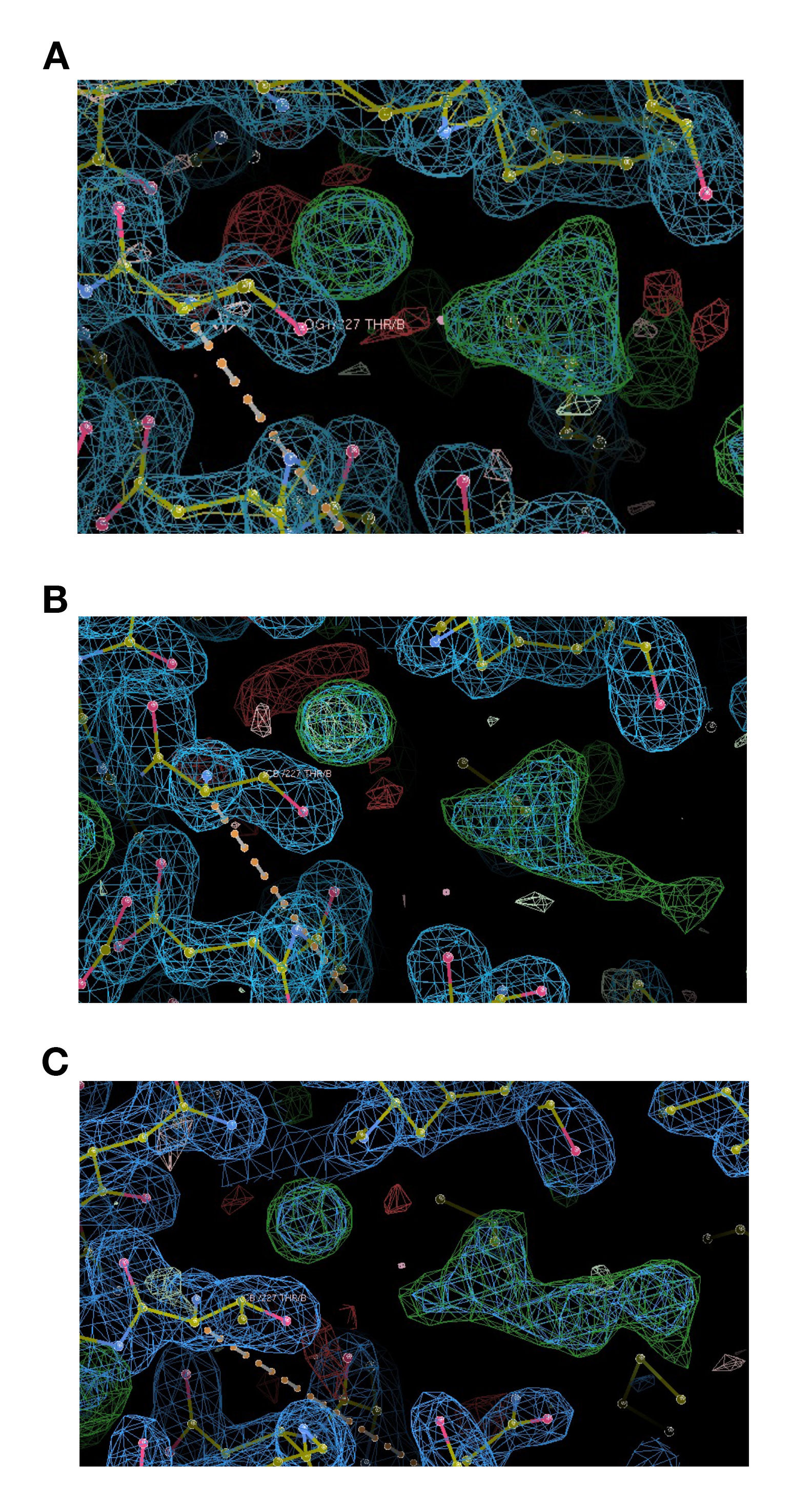

**Figure S5**. Raw data of colorimetric assay for Nyl10 and Nyl12 with **A)** N-(4-nitrophenyl) butanamide (N4NB), **B)** N-(4-aminophenyl) butanamide (N4AB). Assay was carried out in triplicates; no enzyme control is indicated as Control.

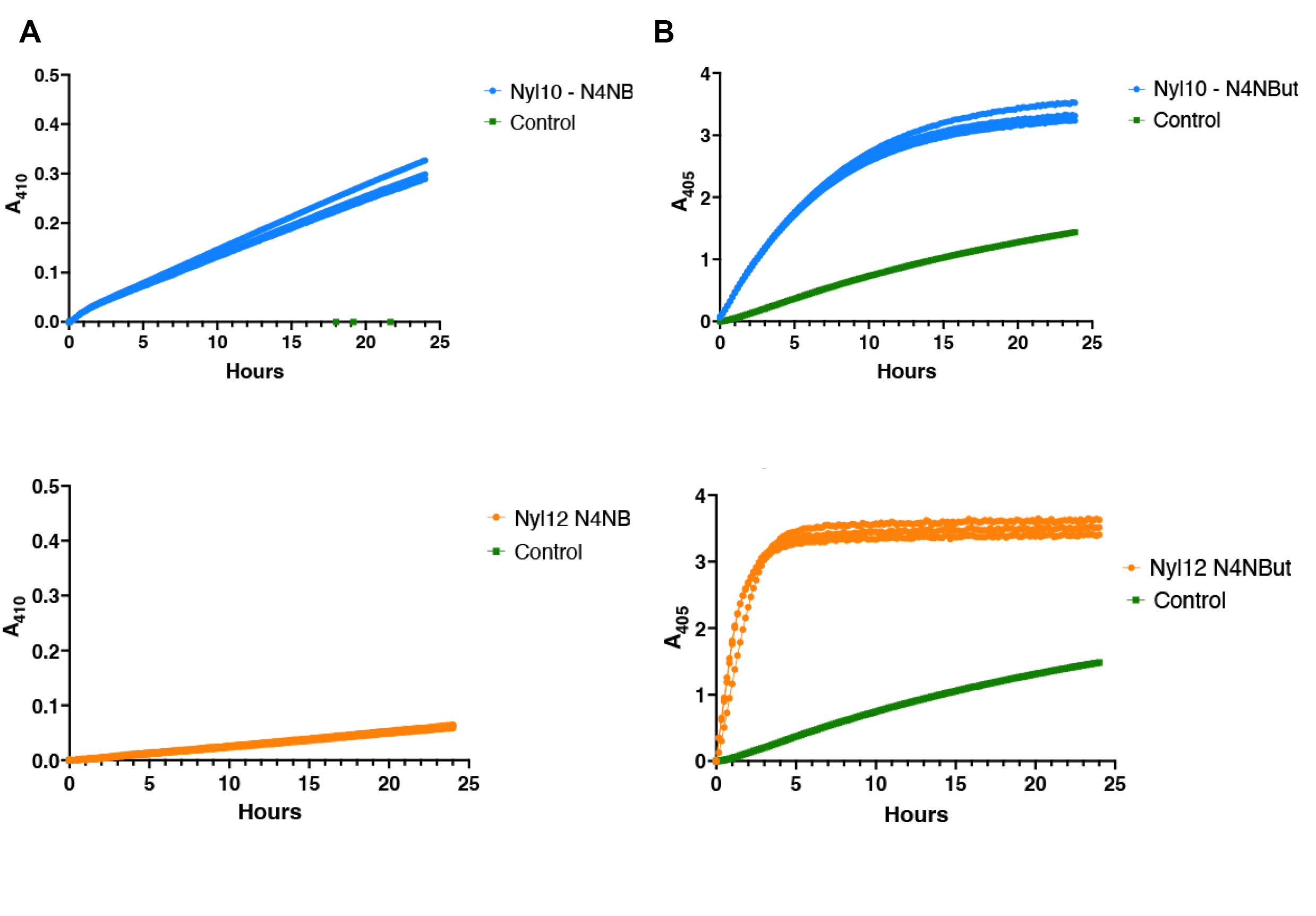

**Figure S6.** Enzyme-substrate interactions in Boltz-2 top model of Nyl12 with capped PA66-L2 substrate. Interacting residues are labeled and shown as sticks. Interactions are shown as dashed lines. Colors: substrate, cyan sticks; chain A, green sticks; chain D, yellow stick; chain C, magenta sticks. Nonpolar interactions, dark blue dashes; hydrogen bonds, orange dashes; Salt bridge (K231), light blue.

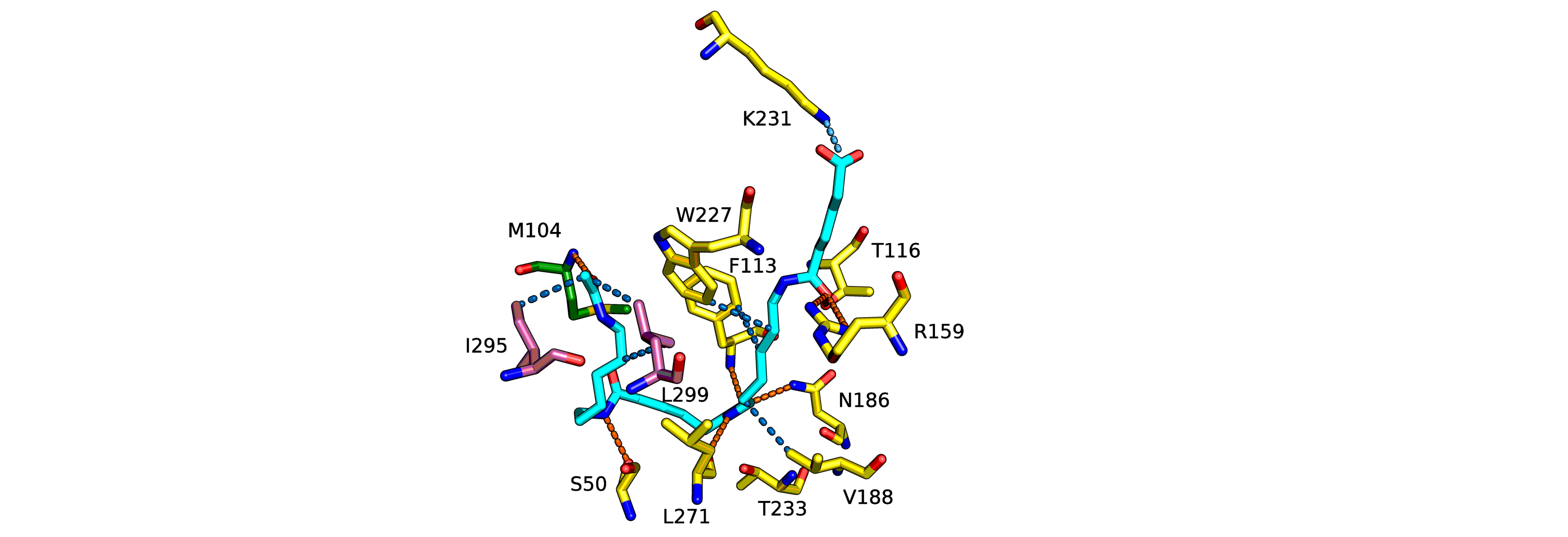

**Figure S7**. DLS profiles of Nyl10, Nyl12 and Nyl50 at ~1 mg/ml.

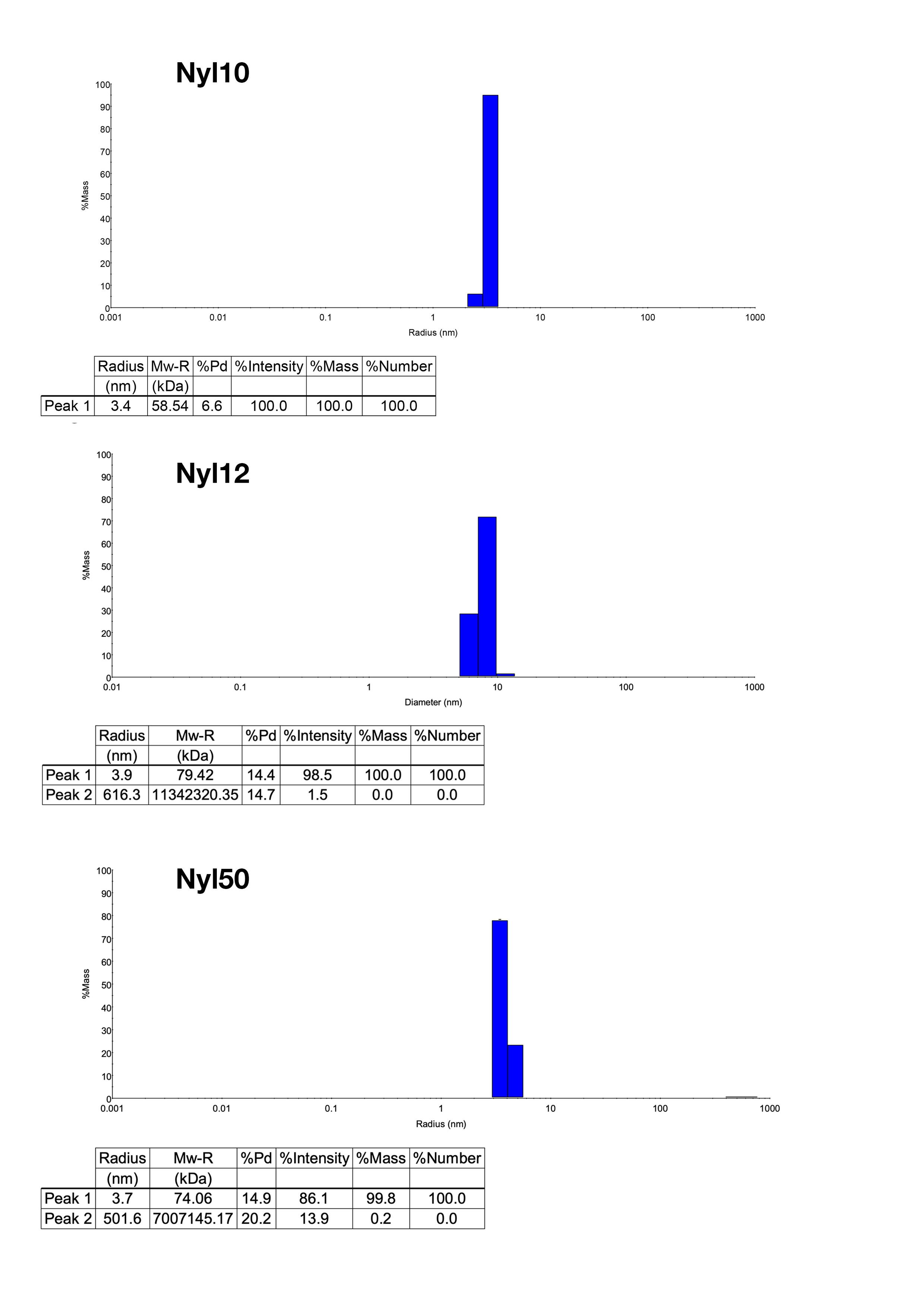

**Figure S8**. **A.** Nyl10 analytical ultracentrifuge coefficient plot at 30, 150 and 300 µM (0.1, 0.5 and 1.0 mg/ml). **B**. Nyl10 analytical ultracentrifuge residue/fit plot for the same concentrations.

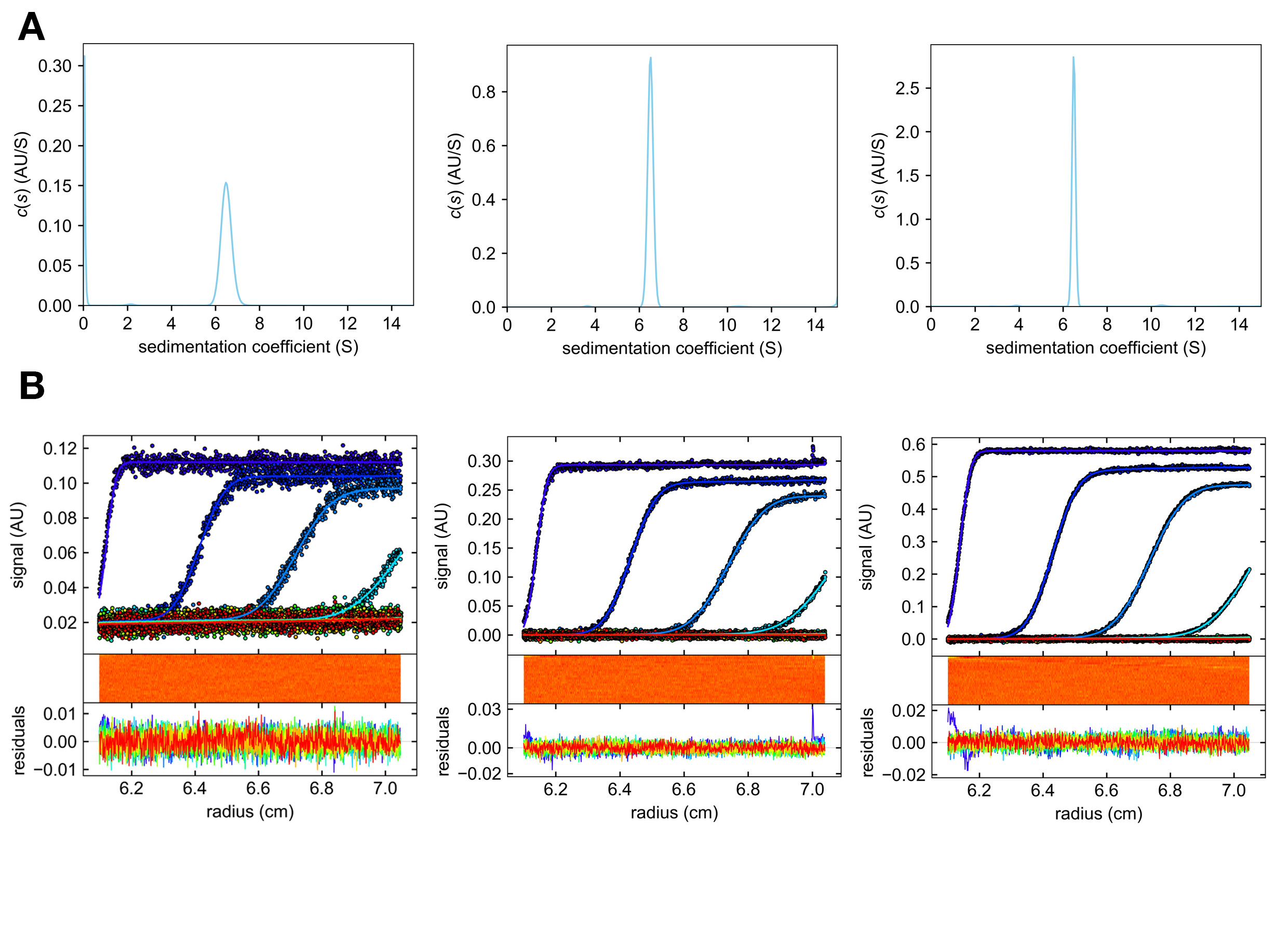

**Figure S9**. **A.** Nyl12 analytical ultracentrifuge coefficient plot at 3, 15 and 30 µM (0.1, 0.5 and 1.0 mg/ml). **B**. Nyl12 analytical ultracentrifuge residue/fit plot for the same concentrations.

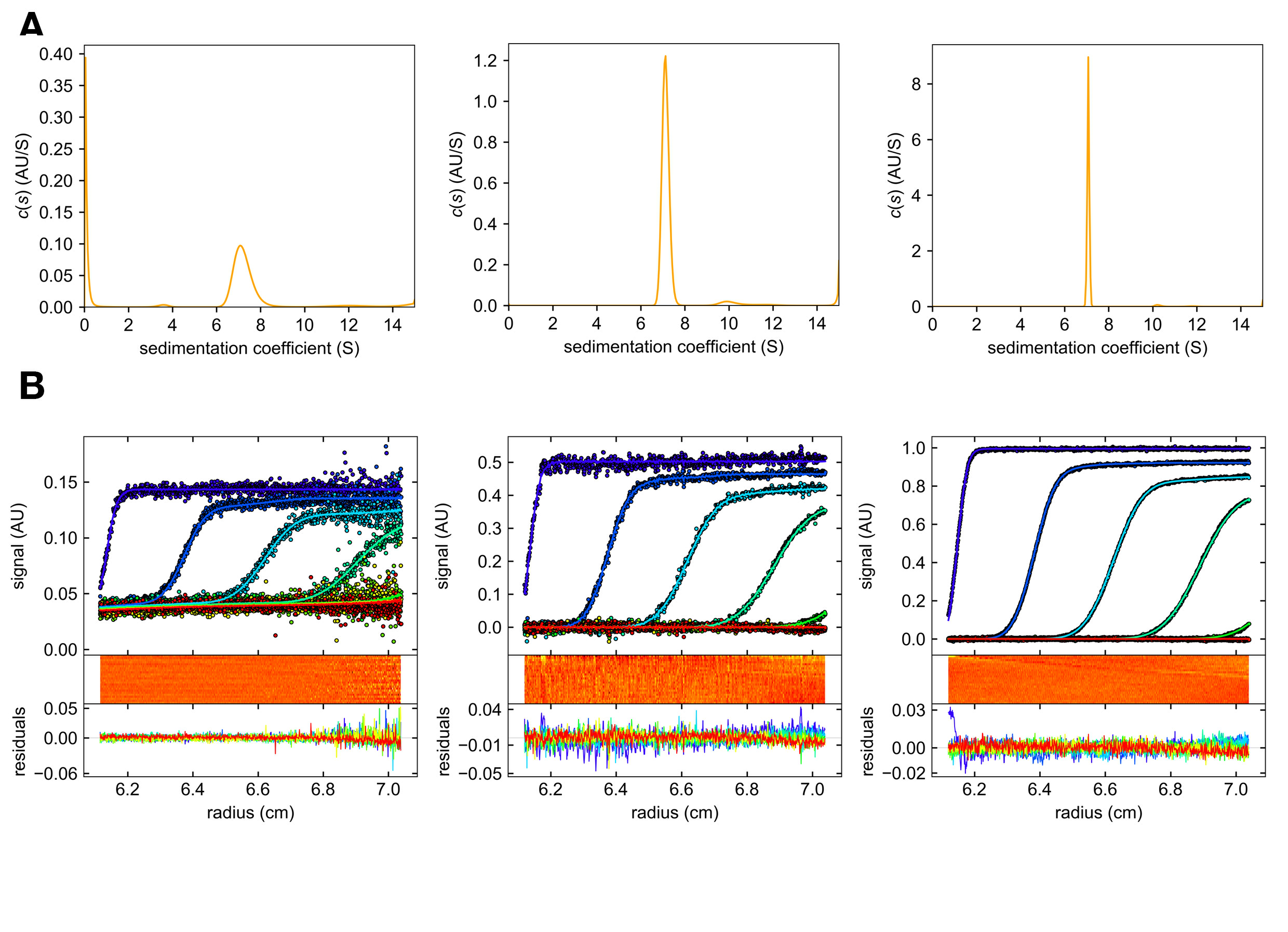

**Figure S10**. **A.** Nyl50 analytical ultracentrifuge coefficient plot at 30, 150 and 300 µM (0.1, 0.5 and 1.0 mg/ml). **B**. Nyl50 analytical ultracentrifuge residue/fit plot for the same concentrations.

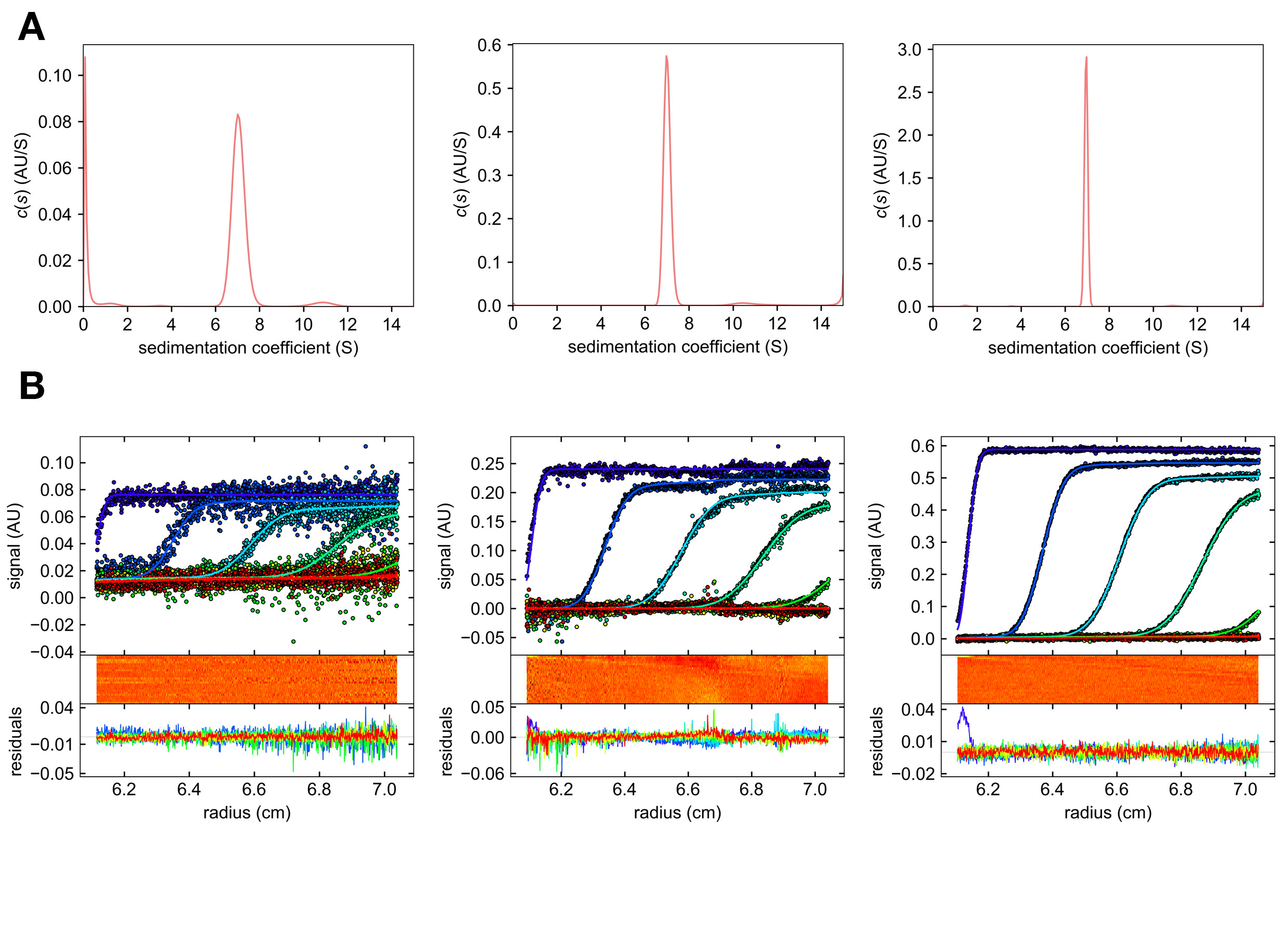

**Figure S11**. **Comparative Efficiency of Nylon Hydrolases toward PA66 and PA6. A-C.** Enzyme-saturated conventional Michaelis–Menten kinetics were determined by measuring reaction rates as the amount of released 6-AHA equivalents in reactions with increasing concentrations of PA66, 500 nM enzyme (75 °C, 200 mM Tris buffer, pH 8, 2 h).

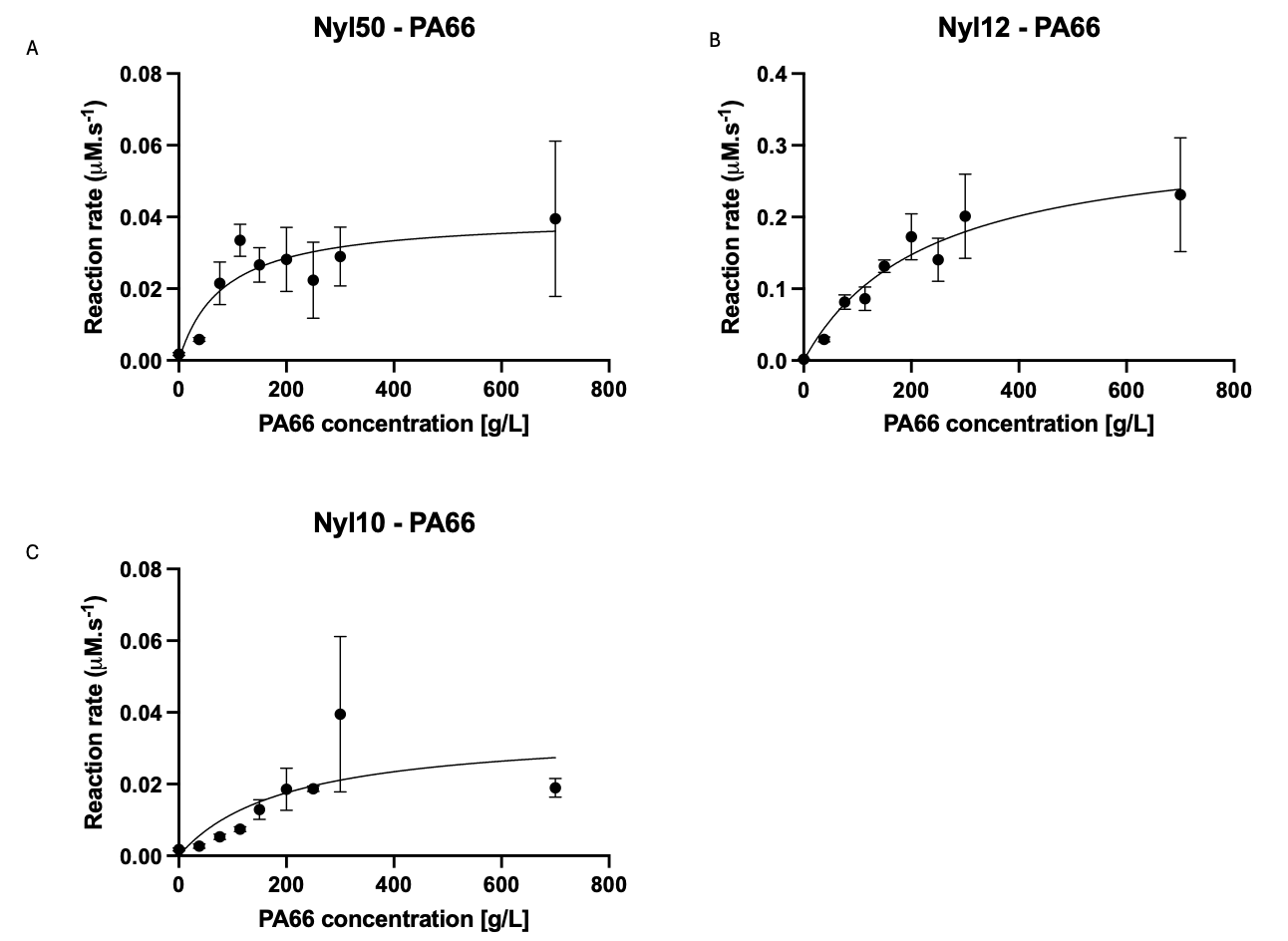
